# Species-specific transformer models of bacterial gene order and content for genomic surveillance tasks

**DOI:** 10.64898/2026.04.28.721069

**Authors:** Samuel T. Horsfield, Maciej Wiatrak, James O. McInerney, Stephen D. Bentley, Caroline Colijn, John A. Lees

## Abstract

Transformer models enable functionally meaningful representation of complex biological data, such as nucleotide or protein sequences. Existing foundation transformer models are trained on large multi-domain corpuses of unlabelled DNA or protein data, showing unmatched task generalisation. However, these foundation models are often outperformed on domain-specific tasks by models trained on taxonomically-constrained data, such as prokaryote gene annotation. By extension, species-specific transformer models hold promise for targeted analyses, given sufficient training data are available. Epidemiological analysis of bacterial pathogens exemplifies the use case of species-specific transformers, due to the wealth of genome data available, coupled with pathogen-specific analyses carried out during routine and outbreak surveillance. Here, we trained a transformer model, PanBART, on the gene content and gene order of two important and biologically distinct bacterial pathogens, *Escherichia coli* and *Streptococcus pneumoniae*, benchmarking against state-of-the-art non-transformer approaches for genomic epidemiology. We show PanBART learns representations of population structure in an unsupervised manner, and can be used to accurately assign genomes to biologically-meaningful sequence clusters. PanBART is also able to identify emergent lineages, differentiating them from pre-existing lineages, and can accurately predict genomes likely to uptake genes involved in antibiotic resistance before a transfer event has occurred. Finally, PanBART can be used to conduct co-selection analysis to identify pairs of genes likely to be evolving together. Our work demonstrates that species-specific transformer models can be employed in many critical public health scenarios. We lay the groundwork for wider application of such models in epidemiological analysis, and provide scenarios where such models excel.

## Introduction

Transformer models, exemplified by large language models (LLMs), enable data-driven feature extraction from sequential data, providing highly flexible frameworks for classification and regression tasks (Devlin et al. 2018; Lewis et al. 2019; Vaswani et al. 2017). These models predict a masked element in a sequence, known as a “token”, based on the identity and position of other tokens present in the sequence. The sequential nature of one-dimensional nucleotide and protein sequences, analogous to natural language, has led to application and multi-domain development of transformer models trained on biological data. Foundation transformer models, such as DNABERT (Ji et al. 2021), Nucleotide Transformer (Dalla-Torre et al. 2025) and ESM (Lin et al. 2023), are trained on vast corpuses of unlabelled biological sequence data from across the tree of life, enabling them unparalleled generalisation capability to a large range of tasks. Nucleotide sequence model applications include gene expression prediction, protein binding and epigenetic modification sites (Ji et al. 2021; Dalla-Torre et al. 2025; Avsec et al. 2021, 2026), while protein sequence models allow for structural prediction, sensitive homologue clustering, functional annotation and mutation effect prediction (Lin et al. 2023; Talo and Bozdag 2025; Brandes et al. 2023).

Due to the wealth of publicly-available microbial genome data from extensive surveillance and sampling efforts (Hunt et al. 2024; Richardson et al. 2023; Schmidt et al. 2024; Timme et al. 2019), some transformer model training has focused exclusively on taxonomically-restricted datasets, such as all the genomic variation in a population, known as a “pangenome” (Tettelin et al. 2005). Given the greater specificity of this model training, these approaches either outperform foundation models for domain-specific tasks, or provide additional functionality not available in a more generally trained model. ProkBERT is a transformer model trained on nucleotide sequences from bacteria, archaea, viruses, and fungi, and is capable of microbial-specific genome feature classification tasks, such as promoter and phage identification (Ligeti et al. 2023). However, ProkBERT is trained on short segments of input genomes, due to computational limitations regarding attention window size, reaching up to 2048 bp for the largest model. In the case of bacteria, a large degree of within-species variation arises at the level of genome architecture between individuals, specifically gene content (orthologous gene presence or absence) and gene order (orthologous gene chromosomal ordering) (McInerney 2026; Brockhurst et al. 2019). Furthermore, non-transformer approaches have shown that the presence or absence of a given gene can be predicted from the presence or absence of other genes in the genome (Beavan et al. 2024; Whelan et al. 2020). Therefore, representations of gene content contain learnable features that can be utilised by a transformer model, as with nucleotide sequence, but in a more compact form, numbering in the thousands of genes rather than the millions of nucleotides per genome. Confirming this hypothesis, a novel transformer model, Bacformer, is trained on only bacterial gene content and order from metagenome data, and therefore captures full genome architecture in its context window (Wiatrak et al. 2025). As Bacformer captures a higher-level representation of genome variation, it outperforms foundation models on gene-specific classification tasks, such as operon identification and gene essentiality prediction. Therefore, transformer models trained on taxonomically-restricted data, including on a gene-level rather than nucleotide-level, hold promise for a range of tasks on which they outperform general foundation models.

As taxonomically-restricted training improves transformer model specificity and functionality, further taxonomic restriction to single species would enable models to be applied to distinct population-specific tasks, such as the genomic epidemiological analysis of bacterial pathogens. In this context, individuals of a species are clustered based on genomic, and likely phenotypic, similarity to produce “lineages” (a group of genomes arising from a single common ancestor) (Croucher and Didelot 2015; Lo et al. 2019; Horesh, Blackwell, et al. 2021; Lees et al. 2019). Researchers can then place newly sequenced genomes within existing or novel lineages, enabling identification of transmission chains and emergence of potential outbreaks (Batisti Biffignandi et al. 2023; McHugh et al. 2025). Transformer models are well-suited for epidemiological analysis of bacterial pathogens, which usually require a suite of pathogen-specific tools to conduct genomic epidemiology (Argimón et al. 2021; Lee et al. 2019; Petit and Read 2020), but could be streamlined to a single specific-specific transformer model. Furthermore, hundreds of thousands of genomes are now available for certain bacterial pathogen species through genomic surveillance efforts (Timme et al. 2019; Hunt et al. 2024), providing sufficient data from transformer model training. However, no previous transformer models have focused solely on training using a single species dataset.

Several distinct transformer architectures could be applied to single-species genome datasets, with choice dictated by the biological relationship being captured. Autoregressive models, such as Nucleotide transformer (Dalla-Torre et al. 2025), learn relationships between tokens in a unidirectional fashion, meaning only upstream tokens are used in the prediction of a masked token. Alternatively, transformer models with a “Masked Language Modelling” (MLM) objective, such as DNABERT (Ji et al. 2021) learn token relationships in a bidirectional fashion, meaning tokens upstream and downstream of a masked token are used in its prediction. As the order in which a genome is read is arbitrary, MLM models are better suited to modelling gene-gene relationships. A hybrid approach, BART (Bidirectional and Auto-Regressive Transformer), combines both an autoregressive and MLM learning objective, and has been shown to be more robust to extreme data permutations than MLM or autoregressive models, such as deletions and sentence reordering (Lewis et al. 2019). A BART-based approach has not been previously applied to genome data; however, due to the arbitrary ordering of contigs in genome assemblies, typical training of transformers on biological data involves random reordering and inversion of contigs (Wiatrak et al. 2025). This data permutation is similar to the extreme data permutations to which BART models are robust to, meaning they are likely well suited to this data type.

In this work we explore the use of single-species transformer models in the context of genomic epidemiological analysis of bacterial pathogens. We train a BART transformer model, which we call “PanBART”, on representations of gene order and gene content from two important bacterial pathogens with distinct population structure and genome architectures, *Streptococcus pneumoniae* and *Escherichia coli*. We apply PanBART to a series of tasks covered by state-of-the-art tools, including genome clustering and lineage assignment, emergence detection, gene uptake prediction and co-selection analysis, and explore the effect of dataset sampling on model performance. We show that PanBART is highly accurate in all tasks it is applied to, often outperforming state-of-the-art methods. Our work is the first demonstration that pathogen-targeted models are applicable to a wide range of epidemiological analysis tasks, providing a case for their widespread use in public health as a direct replacement for existing tools and workflows.

## Results

### Training a transformer model on bacterial gene order and gene content

We developed a reusable workflow for generating representations of bacterial gene order and content, and subsequent training of transformer models. To capture a representative sample of species genomic diversity, we first identified high-quality genomes (**Figure 1A**, **Methods**) in the AllTheBacteria dataset (Hunt et al. 2024). We limited model training and analysis to *Streptococcus pneumoniae* and *Escherichia coli*, which had large numbers of genomes (119,432 and 393,738 genomes used in training/analysis respectively), and differ in their genome biology; *S. pneumoniae* is able to naturally incorporate extracellular DNA into its genome, known as “competence”, possessing a relatively small pangenome in which genes are shared broadly across genetically equidistant lineages (Croucher et al. 2014), while *E. coli* is not naturally competent, and has a large pangenome whereby gene flow is more restricted to closely related lineages (Horesh, Taylor-Brown, et al. 2021).

**Figure 1:**
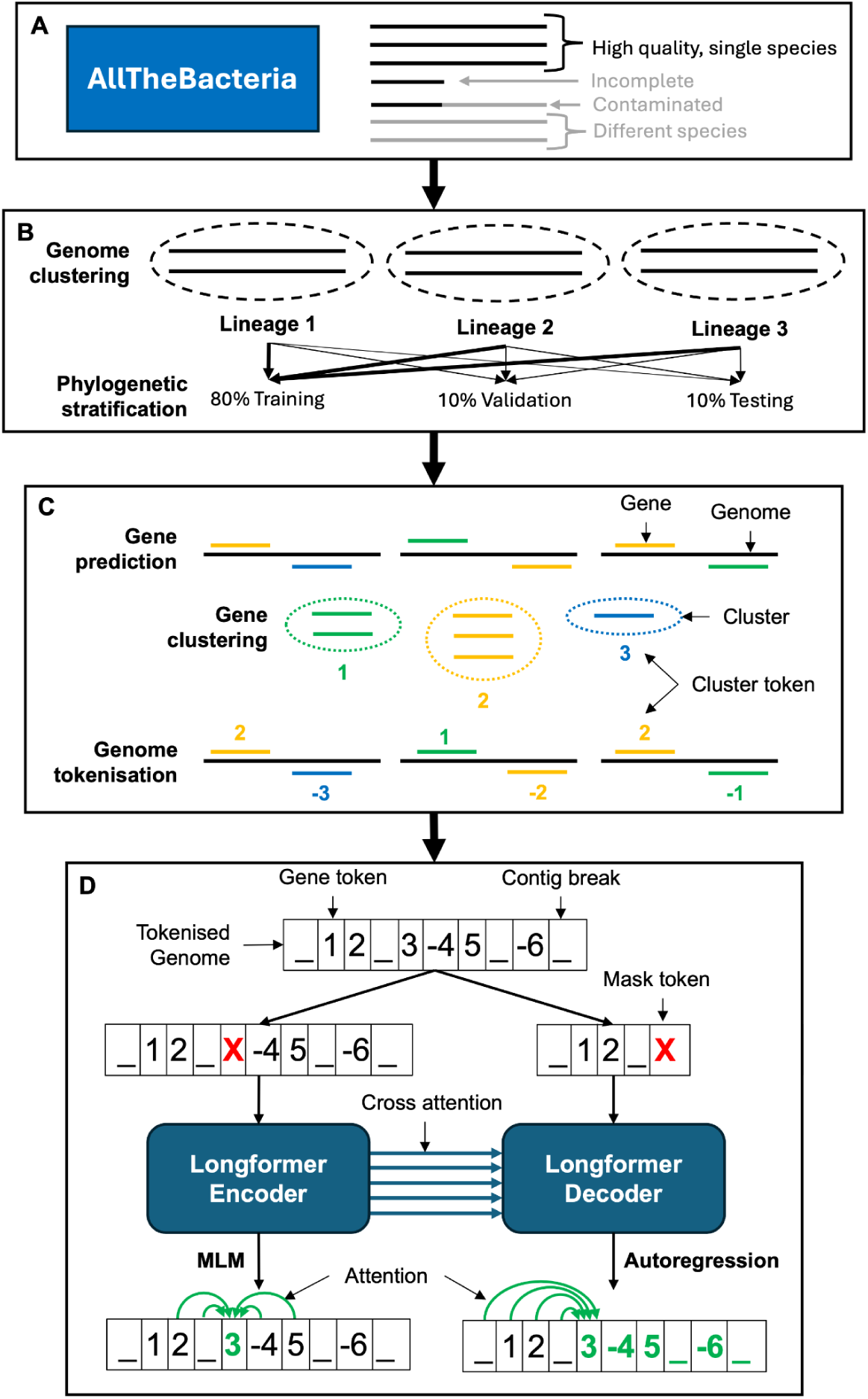
PanBART training overview. (**A**) Genomes for a specific species are sampled from AllTheBacteria using species designations from Sylph. Only high quality genomes are selected (see **Methods**). (**B**) Sampled genomes are clustered into lineage clusters using PopPUNK. Genomes are sampled in a stratified manner, with each PopPUNK lineage cluster being randomly split into training, validation and testing dataset in an 80%-10%-10% split. (**C**) Genes are predicted, clustered based on amino-acid sequence identity, and tokenised in each genome across the full unstratified dataset using WTBCluster (see **Methods**). (**D**) Tokenised genomes are passed through the Longformer encoder-decoder architecture with two training objectives. In the Masked-Language-Modelling (MLM) training objective, mask tokens are added randomly throughout the tokenised genomes and passed through the Longformer encoder, which uses bidirectional attention to infer masked tokens from tokens surrounding the masked token. In the autoregression training objective, downstream tokens are masked, with unidirectional attention used to infer masked tokens using tokens upstream of the masked token. Cross attention between the hidden layers of the encoder and decoder is used to provide contextual information to improve the accuracy of autoregression.

To ensure training data represented the full diversity of each species, we clustered individual species genome datasets using PopPUNK (Lees et al. 2019) before conducting stratified sampling (**Figure 1B**). PopPUNK places closely-related genomes into sequence clusters, referred to here as “lineages”, based on pairwise genome distances. Each lineage was then split into training, validation and testing datasets (80%-10%-10% of each lineage respectively), meaning that all lineages were represented in the training, validation and testing datasets. Lineage sizes ranged from 1-35691 (median=2) genomes and 1-5870 (median=3) in *E. coli* and *S. pneumoniae* respectively (**Supplementary Figure 1**). Genomes in each dataset were then tokenised, whereby each genome is represented as a string of integers, each integer representing a gene cluster (**Figure 1C**). The ordering of integers represents the gene order in each genome, with contig breaks also encoded to ensure the model learns that orientations and positions of contigs are arbitrary.

PanBART was trained on tokenised genomes, using the Longformer-Encoder-Decoder BART architecture (Beltagy et al. 2020) (**Figure 1D**). We chose this architecture as it possesses an increased context window (∼10,000 tokens) compared to standard transformers (∼2,000 tokens), enabling a full large bacterial genome to be used in training. Furthermore, this BART architecture employs both a bidirectional MLM learning objective using an encoder, and unidirectional autoregressive learning objective using a decoder, making this architecture more robust to extreme data permutations than MLM or autoregressive models (Lewis et al. 2019). During training, contig order and orientation were randomised each training cycle, or “epoch” to prevent the model overtraining on a specific arbitrary contig ordering, representing extreme data permutations BART models are robust to. The encoder leverages self-attention only, where only tokens in the sequence fed into the encoder are used during training, while the decoder uses both self-attention and cross-attention with the encoder, meaning that both sequences fed into the encoder and decoder are used for decoder training. Due to the arbitrary ordering of contigs within genomes, we did not include start or stop tokens at the end of sequences, which are normally used in natural language modelling (Vaswani et al. 2017), instead starting and ending each contig with a contig break token.

We trained two separate PanBART models on *S. pneumoniae* and *E. coli* datasets (all genome identifiers available in **Supplementary File 1**, summary of total genomes used in **Supplementary Table 1**). To test the impact of training dataset size and diversity, we independently trained PanBART on 10% and 100% of each dataset, as well as separately holding out a subset of lineages, producing eight trained models per species. Model sizes ranged from ∼28 million to ∼51 million parameters for *S. pneumoniae* and *E. coli* datasets respectively, taking between 190-632 hours for model training depending on the dataset size (**Supplementary Table 2**). These datasets are referred to based on their species, amount of data and the lineages sampled e.g. Pneumo10Sub refers to the dataset of *S. pneumoniae*, containing 10% of all data and a subset of the total lineages (full nomenclature in **Supplementary Table 2**). PanBART was trained on all datasets successfully, shown by reductions in cross-entropy loss coupled with increased per-token prediction accuracy (**Supplementary Figure 2**). Models trained on 100% of the data converged in fewer epochs than those trained on 10% due to the increased volume of data enabling fewer passes through each dataset to reach the same level of accuracy. Final epoch per-token prediction accuracy was high for all models, with precision, recall and F1-score consistently around ∼0.94, with the exception of Pneumo10Sub, which had final-epoch per-token prediction precision, recall and F1 score values of 0.84, 0.84, 0.83 respectively, compared to Pneumo10All, which had values of 0.93, 0.92, 0.92 respectively. Gene frequency was positively associated with PanBART gene prediction confidence, with genome randomisation removing this association, meaning that gene order and gene frequency are crucial features learned by PanBART (**Supplementary Figure 3**).

### Gene order enhances epidemiological clustering over gene content alone

A key practice in genomic epidemiology is the differentiation of genetically similar clusters of genomes to identify potential outbreaks and transmission chains (Croucher and Didelot 2015). Clustering can also be conducted using transformer models, using their internal representations of tokens as N-dimensional float vectors, known as “embeddings” (Wiatrak et al. 2025; Lin et al. 2023). The distance between embeddings represents their semantic similarity; in the case of genes, grouping clusters with similar embeddings generates clusters of functionally similar genes. Furthermore, averaging gene-level embeddings across an entire genome, known as “mean-pooling”, can be used to generate genome-level embeddings, which can then be clustered into groups of closely-related genomes. Although genome-level embeddings have been used to cluster genomes at a species level (Ligeti et al. 2023; Wiatrak et al. 2025), the ability to cluster at lineage-level classification schemes necessary for genomic epidemiology has not been investigated. Gene content alone is not sufficient to generate phylogenetically consistent clusters (Lees et al. 2019, 2018); therefore we investigated the effect of combining gene content and gene order, learned by PanBART, on the ability to cluster genomes at below-species resolution. If it is the case that closely-related genomes have closer embeddings than those of distinct lineages, the representation of genome architecture learned by PanBART can be used to cluster genomes into lineages for epidemiological purposes.

To test the ability of PanBART to generate epidemiologically-informative embeddings, we passed genomes used in training and those held out for testing through each PanBART model, and generated genome-level embeddings with a total of 256 dimensions, as described above. We compared PanBART to accessory distances calculated from Sketchlib (Lees et al. 2022), which uses only gene presence/absence to cluster genomes, generating embeddings from the pairwise distance matrix generated by Sketchlib. Sketchlib is implemented in PopPUNK to calculate pairwise genome distances (Lees et al. 2019), and therefore represents the state-of-the-art for genomic epidemiology analysis. Furthermore, Sketchlib is highly scalable, enabling analysis of 100,000s genomes, which is not possible with other existing genomic epidemiology tools. To compare the tools, we used per-cluster Silhouette score, which measures how closely samples with the same ground-truth label lie within N-dimensional space. A higher Silhouette score indicates that samples which share a label tend to cluster together, while a lower score means samples with the same label are more dispersed in embedding space. We used the original PopPUNK lineage assignments as ground-truth labels, which uses information on gene presence/absence and core mutations for lineage assignment, calculating Silhouette scores using the original embedding dimensions of each method, and using UMAP for visualisation only.

For almost all models and species combinations, PanBART (gene content + order) had higher median Silhouette scores than Sketchlib (gene content only) for the 10 largest lineages, meaning that PanBART embeddings have a better agreement with PopPUNK clusters (gene presence + core mutations) than those of Sketchlib (**Figure 2**). An exception was the Pneumo10All model, which had a lower Silhouette value than the corresponding Sketchlib analysis (**Supplementary Figure 4**). PanBART clusters were also visually more coherent than those of Sketchlib, with PanBART embeddings forming larger single-colour clusters compared to those of Sketchlib. This same performance improvement in PanBART was observed in testing data (**Supplementary Figure 5**), as well as models trained on a subset of total lineages (**Supplementary Figures 6 & 7**). Finally, PanBART outperformed Sketchlib when clustering genomes from lineages not included in training data, producing embeddings with higher Silhouette score than Sketchlib (**Supplementary Figure 8**). Therefore, PanBART generalises to clustering genomes from lineages not previously seen in training, known as “zero-shot classification”, indicating that it is not overtrained, and can therefore be applied to previously unseen data.

**Figure 2:**
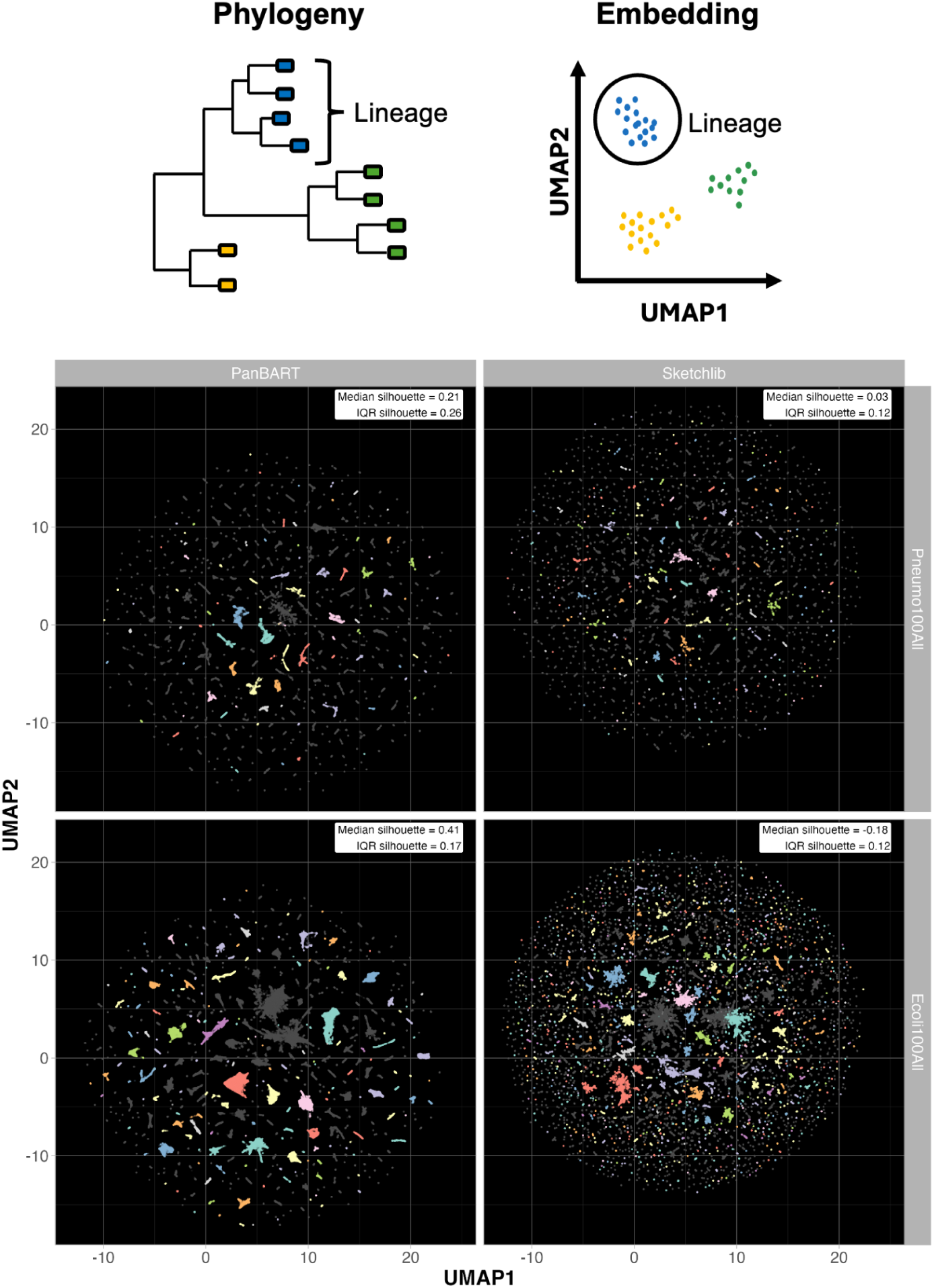
Representation of phylogenetic information in embedding space. Genomes belonging to the same lineages should be placed in the same region of embedding space (*top*). Comparison of PanBART and Sketchlib genome embeddings (*bottom*). Each point represents the embedding of a single genome. Dataset names displayed on vertical facets (dataset labelling nomenclature is available in **Supplementary Table 2**). Embeddings are shown for genomes used in training datasets only. Colours show the 10 largest PopPUNK lineage clusters for each species. The median and Inter-Quartile Range (IQR) of silhouette scores for the 10 largest PopPUNK lineage clusters is shown in the top right corner of each plot.

A key functionality of any genomic epidemiological tool is the ability to assign newly-sequenced query genomes to existing clusters. **Figure 2** highlights that PanBART embeddings represent population structure, meaning distances between genomes in embedding space represents genome similarity. Therefore, we hypothesised that query genomes could be assigned to existing clusters by measuring embedding distances between query and existing genomes with known lineage labels, and assigning query genomes to the lineage they are closest to. To benchmark per-genome lineage assignment, we provided genomes held out of training to the model, mimicking the process of ongoing genome surveillance, whereby previously unseen genomes are classified to existing lineages. We then assigned PopPUNK lineage labels to these genomes using k-Nearest Neighbours (kNN) classification based on the full embeddings (not 2D UMAP) from PanBART or Sketchlib, using k varying from 1 to 100. kNN measures the embedding distances between a query genome with other labelled genomes, and assigns a label to the query with which the majority of the k-nearest neighbours share. By increasing k, more labelled data is considered in label assignment, reducing sensitivity to outliers at a cost to reduced sensitivity to local patterns, causing underfitting. As the PopPUNK lineage designations of the query genomes were known *a priori*, we determined the accuracy of cluster assignment by comparing predicted and ground-truth labels for both *S. pneumoniae* and *E. coli*.

Overall, Sketchlib performed best for lineage assignment, although differences in performance between PanBART and Sketchlib varied by species and dataset size (**Figure 3**). For both PanBART and Sketchlib, lineage assignment accuracy was highest at k=1 for both species. PanBART was also able to assign genomes at ∼170 and ∼112 genomes per minute for *S. pneumoniae* and *E. coli* respectively, which did not depend on dataset size (**Supplementary Table 3**). Sketchlib was ∼10x faster than PanBART, with sketching and distance calculation rates of ∼2210 and ∼1200 genomes per minute for *S. pneumoniae* and *E. coli* respectively, which was also independent of dataset size.

**Figure 3:**
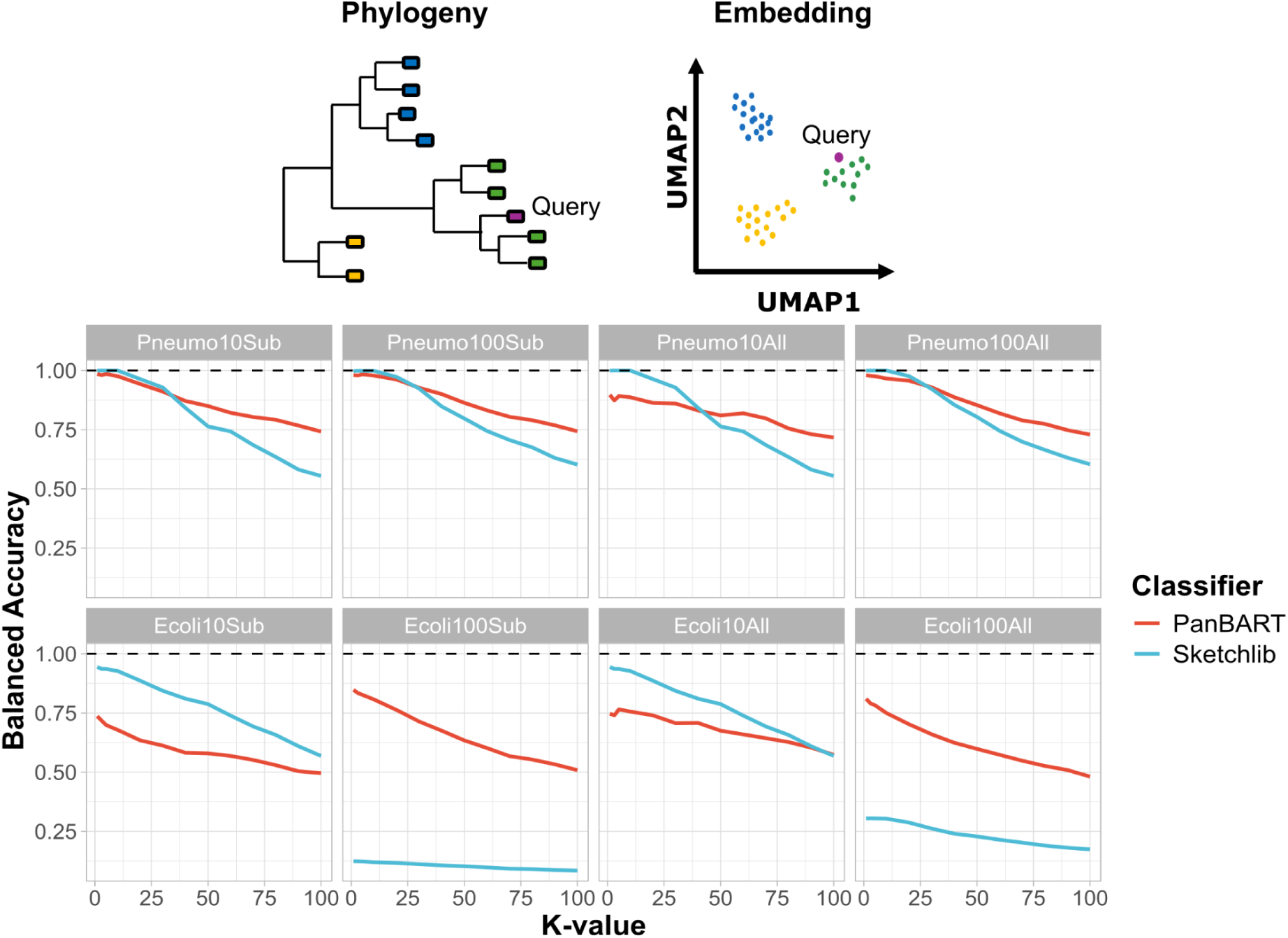
Lineage classification accuracy of query genomes. Query genomes closely related to genomes from known lineages should be placed close in embedding space and assigned the same label (*top*). Comparison of lineage classification accuracy between PanBART and Sketchlib (*bottom*). Each line represents the balanced accuracy of PopPUNK lineage cluster assignment based on k-nearest neighbours (kNNs) in embedding space. PanBART results are shown by red lines; Sketchlib by blue lines. k-values used for classification are shown on the x-axis. Dataset names provided in facet labels (dataset nomenclature available in **Supplementary Table 2**).

For *S. pneumoniae*, Sketchlib accuracy was 100% at k=1, regardless of how the dataset was subsetted. PanBART accuracy was slightly lower than Sketchlib, and was highest for models trained on 100% of the dataset, achieving 98% accuracy at k=1 for both Pneumo100All and Pneumo100Sub models. For *E. coli*, Sketchlib again performed best, achieving 94% accuracy at k=1 on the Ecoli10All and Ecoli10Sub datasets. PanBART *E. coli* performance was worse than with *S. pneumoniae*; models trained on 100% of the dataset again performed best, achieving 81% and 85% accuracy at k=1 for Ecoli100All and Ecoli100Sub models respectively.

By increasing k, we were able to confirm our observation above that PanBART clusters appear more coherent with true cluster labels than Sketchlib. For all datasets and methods, balanced accuracy fell as k increased, due to inclusion of more differently labelled genomes during classification. For *S. pneumoniae*, Sketchlib slightly outperformed PanBART when k < 25, with all models achieving >0.95 balanced accuracy. An exception to this was the Pneumo10All PanBART model, which had notably lower accuracy at k <50, mirroring its lowered embedding Silhouette values (**Supplementary Figure 4**). At k > 25, PanBART outperformed Sketchlib for all datasets, in line with observations in **Figure 2**, where PanBART forms larger consistently-labelled clusters, which contribute to better performance at large k values.

Performance on *E. coli* classification was more varied between PanBART and Sketchlib. PanBART classification accuracy was consistent across all models, and while Sketchlib outperformed PanBART on the Ecol10Sub and Ecoli10All analyses, Sketchlib performance declined notably on Ecoli100All and Ecoli100Sub datasets. The reason for this performance drop can be observed in **Supplementary Figure 6** and **Figure 2**, where the embedding space for Sketchlib for Ecoli100Sub and Ecoli100All and data is highly unstructured in comparison to Ecoli10Sub and Ecoli10All data. Therefore, the abundance of differently labelled genomes in close proximity in Sketchlib embedding space leads to incorrect query assignment, even when using k=1.

We then investigated whether lineage size impacts accuracy of classification of novel genomes. We found that in both species and all dataset sizes, genomes were more accurately assigned to larger lineages for both PanBART and Sketchlib (**Supplementary Figures 9 & 10**). However, PanBART classification accuracy was comparable to or better than Sketchlib, particularly at higher values of k. Increasing k reduced accuracy of small cluster assignment, likely due to clusters with size < k being merged with larger clusters during classification, resulting in incorrect assignment. Testing zero-shot classification performance, PanBART accurately assigned lineages not included in training, outperforming Sketchlib, and is not impacted by the presence of unknown tokens which do not appear in training data (**Supplementary Figure 11**).

As we observed high consistency between PanBART and PopPUNK lineage assignments, we then tested whether PanBART embeddings could themselves be used to generate epidemiologically meaningful cluster labels. We used Leiden clustering to identify clusters in embedding space, and compared label assignments to those from PopPUNK using Adjusted Rand Index (ARI) and Adjusted Mutual Information (AMI), again comparing performance against Sketchlib. ARI and AMI measure the similarity of genome clustering between two different schemes, with higher values indicating closer agreement. ARI and AMI differ in their scoring of class balancing, with ARI better capturing similarity between equally sized clusters, while AMI is better for clustering schemes with varying cluster sizes. PanBART performed equivalently to Sketchlib, with AMI values >0.7 for both methods across training and testing datasets (**Supplementary Figures 12 & 13**). Therefore, including gene order as well as gene content information did not improve *de novo* inference of meaningful sequence clusters, and may be improved by providing additional phylogenetically informative information, such as using core genome distances as in PopPUNK.

### PanBART can identify emergent lineages using pseudolikelihoods

Transformer models can be highly sensitive to data that is very different from that seen during training, known as “out-of-distribution” data, impacting the ability of models to generalise (Shi et al. 2025). However, this sensitivity can also be leveraged to identify previously unseen data; for example, in predicting deleterious effects of unobserved mutations by mutating all residues *in silico* and measuring the model’s confidence that it would be able to generate such a sequence, known as a “pseudolikelihood” (Meier et al. 2021; Brandes et al. 2023). For epidemiological purposes, a genome that is very different from those previously observed may represent a novel emergent lineage which has a fitness advantage (Balasubramanian et al. 2022). In the case of PanBART, a pseudolikelihood can be calculated for each gene in a given position, and can be summed across a genome to generate a per-genome pseudolikelihood. A per-genome pseudolikelihood is a measure of model confidence that a similar genome was seen during training, with a low pseudolikelihood indicating that a genome is different to training data, and is therefore a potential emergent lineage.

We investigated the sensitivity of PanBART to out-of-distribution data, represented by *S. pneumoniae* genomes belonging to lineages held out of model training (see **Supplementary File 1**). We chose *S. pneumoniae* as its population is made up of equidistant lineages (Croucher et al. 2014), meaning all held-out lineages are equally as distant to all training lineages. We observed no significant difference between training and testing genomes in log pseudolikelihoods, which all belong to lineages used in model training (**Figure 4**). Genomes belonging to “novel” previously unseen lineages had significantly lower log pseudolikelihoods than those from testing data, indicating that PanBART can be used to accurately identify novel emergent lineages as out-of-distribution data. However, despite never observing these novel genomes during training, PanBART was still able to distinguish them from those of *Streptococcus mitis*, a closely related species with which *S. pneumoniae* shares a large degree of gene content (D’Aeth et al. 2021). Genomes held out of training and from *S. mitis* clustered mostly together and independently of other genomes in embedding space (**Supplementary Figures 14 & 15**), although individual lineages can still be differentiated from each other (**Supplementary Figures 8 & 11**). Randomising gene order resulted in the greatest drop in pseudolikelihood, indicating that PanBART is able to detect similarity in gene order across closely-related *Streptococci* (**Supplementary Figure 16**). Results were consistent across models trained on the Pneumo10Sub and Pneumo100Sub datasets, meaning that emergence detection is robust to dataset size. Therefore, PanBART is sensitive to out-of-distribution data, which provides the advantage that genomes from previously observed and unseen lineages can be distinguished.

**Figure 4:**
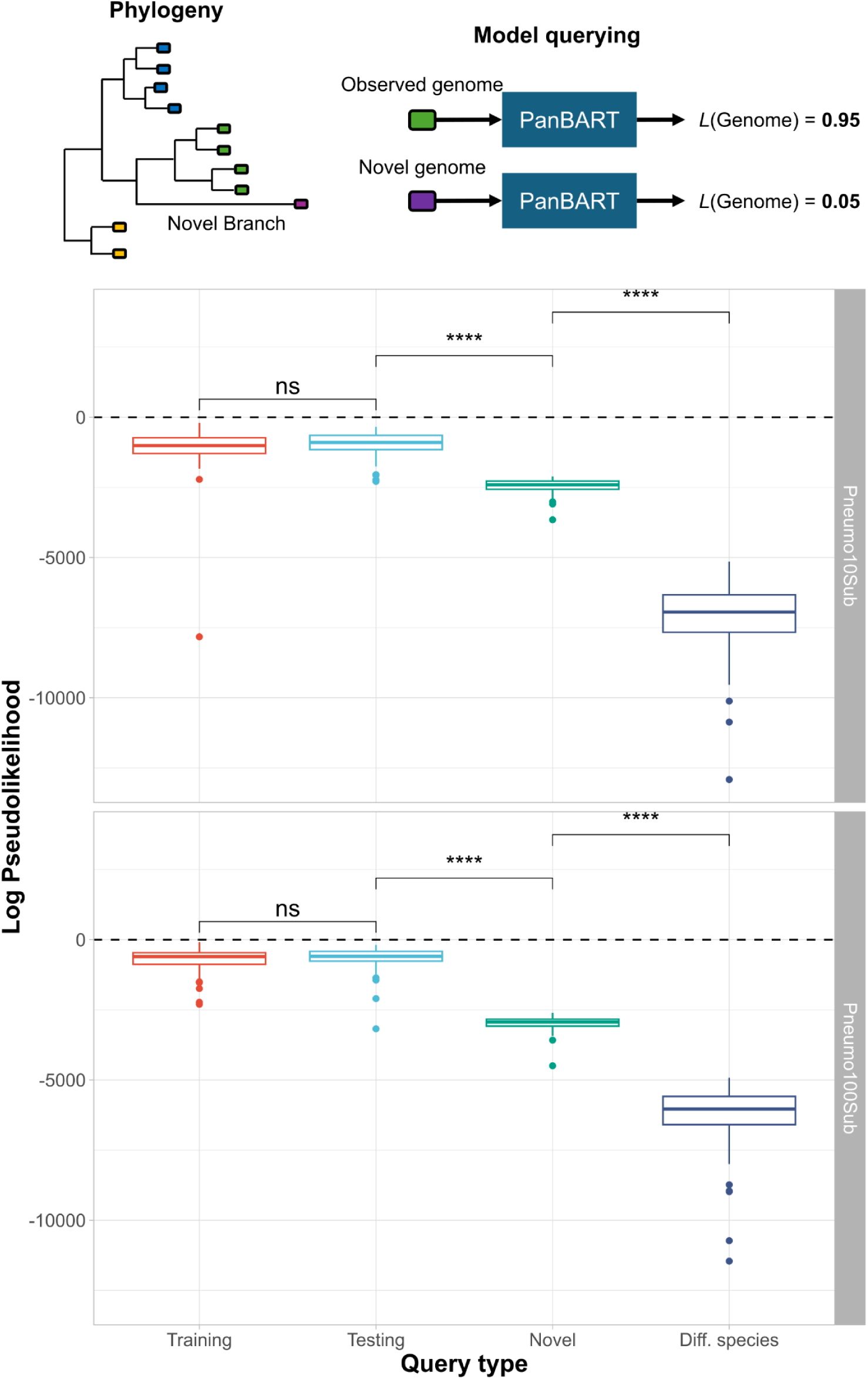
Differentiation of observed and novel lineages by PanBART. Novel genomes belonging to previously unseen lineages should generate low pseudolikelihoods when passed through the model (*top*). Pseudolikelihoods of genomes included and held-out of PanBART training (*bottom*). Query types: “Training”, genomes seen by the model in training; “Testing”, genomes belonging to PopPUNK lineages used in training, but the genomes themselves were not seen in training; “Novel”, genomes belonging to PopPUNK lineages not used in training; “Diff. species”, genomes belonging to a closely related Streptococcal species, *Streptococcus mitis*, which was not used in training. Each query type contains 50 genomes. Statistical comparisons conducted with Wilcoxon test: ns, p >0.05; ****, p ≤ 0.0001.

### PanBART can predict gene uptake potential

Conventional machine learning methods, which do not utilise transformers, have shown great promise in prediction of phenotypic traits from genomic data alone (Biffignandi et al. 2024; Lees et al. 2020; Beavan et al. 2024). These methods tend to identify a small number of predictive features, and ignore higher-order effects, such as gene-gene interactions. Therefore, such methods cannot be used to predict how the genetic background, made up of all loci in a genome, impacts a trait. Transformer models are unique among statistical methods in that they intrinsically learn all second-order interactions (Tomaz da Silva et al. 2025), and so can be applied to cases where a large number of interacting loci across a genome, rather than a small number of independent predictive loci, impact a trait. One such application is the prediction of gene uptake potential, a trait which is dependent on a complex interaction between the availability of external DNA uptake machinery and the symbiotic and antagonistic relationships between genes in a cell (Mell and Redfield 2014; Beavan et al. 2024). Predicting gene uptake potential is of particular epidemiological importance when applied to genes involved in AMR or hypervirulence, with many bacterial pathogen populations harboring lineages with increased drug resistance and virulence (Gladstone et al. 2021; Lo et al. 2022). Transformer models stand as the only existing means of predicting gene uptake potential, providing a novel approach for identifying concerning genome variants before they gain genes that make them more virulent.

We implemented an *in-silico* mutational screen to measure gene uptake potential using individual gene pseudolikelihoods. The screen incrementally adds a gene of interest at all positions in a genome, and calculates the pseudolikelihood of the gene at each position, which describes a model-derived probability of the gene appearing at that location, given the genetic background. We report the highest pseudolikehood value across the genome as a proxy for gene uptake potential prediction. We applied PanBART to predicting uptake potential of *blaCTX-M,* an extended spectrum beta lactamase conferring resistance to beta-lactam antibiotics in *E. coli (Ramadan et al. 2019)*. We chose *blaCTX-M* as it is sufficient for conferring extended beta-lactam resistance, representing a simple single-gene target for testing. Furthermore, *blaCTX-M* presence is strongly associated with specific genetic backgrounds in *E. coli*, meaning we can test whether PanBART is able to associate different genetic backgrounds with *blaCTX-M* presence or absence (Gladstone et al. 2021).

We stratified the *E. coli* dataset into distinct lineages, and measured the association between these lineages and *blaCTX-M*. We included positive controls (labelled “+ve”), in which *blaCTX-M* was known to be present but was removed *in silico*, comparing them to negative samples (labelled “-ve”), in which *blaCTX-M* was naturally absent. We tested the association between *blaCTX-M* with genomes from four lineages: ST131 A and ST131 B/C, part of the high prevalent multidrug resistant ST131 lineage, and ST73 and ST95, two drug susceptible but also highly prevalent lineages (Kallonen et al. 2017; Gladstone et al. 2021). The proportion of genomes containing *blaCTX-M* varied largely between lineages in the dataset spanning from ∼70% positivity in ST131 lineages, to ∼5% in ST73 and ST95 (**Supplementary Table 4**). We would therefore expect pseudolikelihoods to be high for ST131 genomes, and low for ST73 and ST95. Furthermore, each lineage was well represented, with thousands of genomes present in the dataset during training.

We statistically compared all lineages to ST131 B/C (+ve) as a positive control, as we hypothesised these genomes to have the highest overall log pseudolikelihoods due to ST131 B/C having the highest proportion of *blaCTX-M* +ve individuals (**Supplementary Table 4**). Log pseudolikelihoods were not significantly different for ST95 (+ve) and ST73 (+ve) from ST131 B/C (+ve) (**Figure 5A**). Importantly, this result indicates that PanBART is able to positively identify these ST95 and ST73 genomes as having an elevated chance of *blaCTX-M* uptake, despite belonging to lineages with lower *blaCTX-M* frequency. For genomes not naturally containing *blaCTX-M*, there was a significant difference between ST131 B/C (+ve) and ST95 (-ve), and ST131 B/C (+ve) and ST73 (-ve), while no significant difference was identified for ST131 B/C (-ve) and ST131 A (-ve) (**Figure 5B**). This pattern was also observed in training data (**Supplementary Figure 17**). Therefore, PanBART is able to differentiate genomes which do not naturally possess *blaCTX-M*, aligning with their membership of high or low risk lineages.

**Figure 5:**
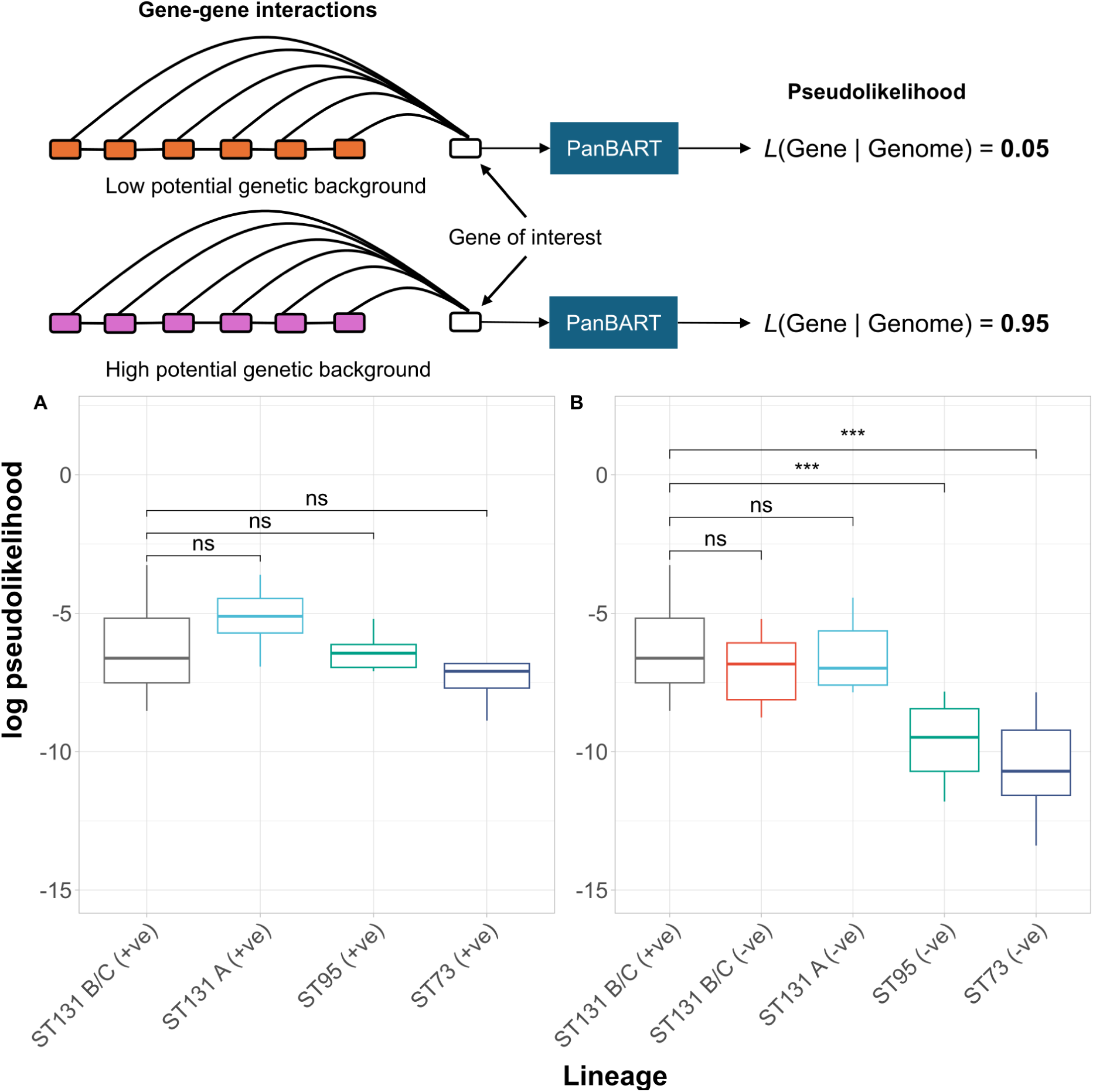
Predicting uptake potential using PanBART. Genetic backgrounds with high association with a gene of interest will have a larger pseudolikelihood (*top*). Log pseudolikelihood comparison of *blaCTX-M* uptake potential across *E. coli* lineages (*bottom*). Analysis was split between genomes which contained *blaCTX-M* but had it bioinformatically removed prior to analysis (labelled “+ve”) (**A**) and genomes which did not contain *blaCTX-M* (labelled “-ve”) (**B**). Comparisons conducted using genomes used in PanBART testing. Each boxplot represents 10 genomes. ST131 B/C (+ve) was shared between both analyses as a positive control for statistical comparisons. Statistical comparisons conducted with Wilcoxon test: ns, p >0.05; *, p ≤ 0.05; ***.

As PanBART can accurately distinguish lineages with high and low risk of acquiring *blaCTX-M*, we next investigated whether PanBART could be used to identify the likely insertion site of *blaCTX-M*. Results showed that for most genomes, the estimate insertion site, given by the locus with the highest pseudolikelihood for *blaCTX-M*, is centred around the true gene location across the genomes analysed, although location estimation is more variable in genomes not included in training (**Supplementary Figure 18**).

### PanBART can detect novel gene co-selection

Epistasis, or co-selection, has been hypothesised to be a driving force for population structure maintenance and speciation due to lowered fitness genotypes missing interacting genes or alleles, known as “outbreeding depression” (Beavan et al. 2023; Neher and Shraiman 2009; Cummins et al. 2026). Identifying interacting loci could therefore be used to predict how populations will evolve by identifying co-evolving genes, and differentiating them from flexibly exchanged genes.

A number of methods have been developed for detecting co-selection, which rely on observing allele co-occurance in a population and using this as a proxy for allele-allele dependencies (Pensar et al. 2019; Whelan et al. 2020; Beavan et al. 2024). Transformer models also learn dependencies between loci, which could act as proxies for potential epistatic interactions (Tomaz da Silva et al. 2025; McInerney 2026). In contrast to existing co-selection detection methods, transformer models can be used to detect co-selection in individual genomes, and so are potentially more sensitive than population-level approaches. Furthermore, transformer models uniquely take into account genome positional information when learning locus associations, which can impact gene expression and function (Hu et al. 2025), and may provide a more accurate prediction as to whether two genes could viably interact to impact fitness.

To investigate whether PanBART could be used for detecting potential epistatic interactions, we calculated SHAP (SHapley Additive exPlanations) values for all loci in a genome with a particular locus of interest. SHAP methods measure feature importance by ablating features from a model and determine the impact on model predictive performance. Removing features that are important in prediction will have a larger negative impact on performance than removing those that have little importance. As no means of accounting for population structure currently exists when training transformer models, we did not remove phylogenetic effects from model predictions.

We used SHAP values to quantify the impact on PanBART prediction of a target locus when removing or adding other loci across the genome. We hypothesise that removing a locus that is under co-selection with a target locus will have a large negative impact on the ability of PanBART to predict the target locus, as these two loci are predictive of each other’s presence or absence. As this is computationally expensive, it was infeasible to do this for all-versus-all genes; therefore we investigated a single target we hypothesised to be under co-selection. Our gene of choice must be in the accessory genome as core genes, such as penicillin binding proteins in *S. pneumoniae* which are known to have co-selected alleles (Skwark et al. 2017), would not have gene presence/absence variations required for detection of gene-gene co-selection. We chose the *cblB* gene in *E. coli*, a gene encoded on an accessory gene cassette involved in the production of the bacteriocin, Colibactin. Colibactin production is tightly regulated by iron homeostasis, with increases in extracellular iron concentration resulting in decrease of colibactin production (Martin et al. 2013), and Colibactin has already been identified to have an antagonistic relationship with antibiotic resistance genes (Mäklin et al. 2025). Therefore, we postulated that *cblB* would be in epistasis with genes involved in iron uptake and metabolism.

We ran PanBART on a selection of *E. coli* genomes stratified from two different lineages across training and testing datasets (**Supplementary File 2**), calculating SHAP distributions across each genome with *cblB* as the target token. We identified a single genome with a SHAP outlier annotated as an aerobactin synthesis locus (Peak 4, **Figure 6A**), which produces a siderophore previously associated to play a role in colibactin regulation (Martin et al. 2013). We were unable to identify the same association when using Spydrpick, a tool which uses mutual information to identify evidence of epistasis in a population-wide manner (Pensar et al. 2019) (**Figure 6B**). Therefore, we believe PanBART is more sensitive than Spydrpick when detecting associations with smaller effect sizes as it can be run on a single genome basis. However, as we could only identify this association in a single genome, PanBART performance is highly dependent on the target genome being analysed. We were also unable to detect evidence of antagonism with antibiotic resistance genes in any tested genome. As we did not account for population structure, many associations may be spurious, leading to a large amount of noise in SHAP distributions that likely confounded identification of co-selection in more genomes.

**Figure 6:**
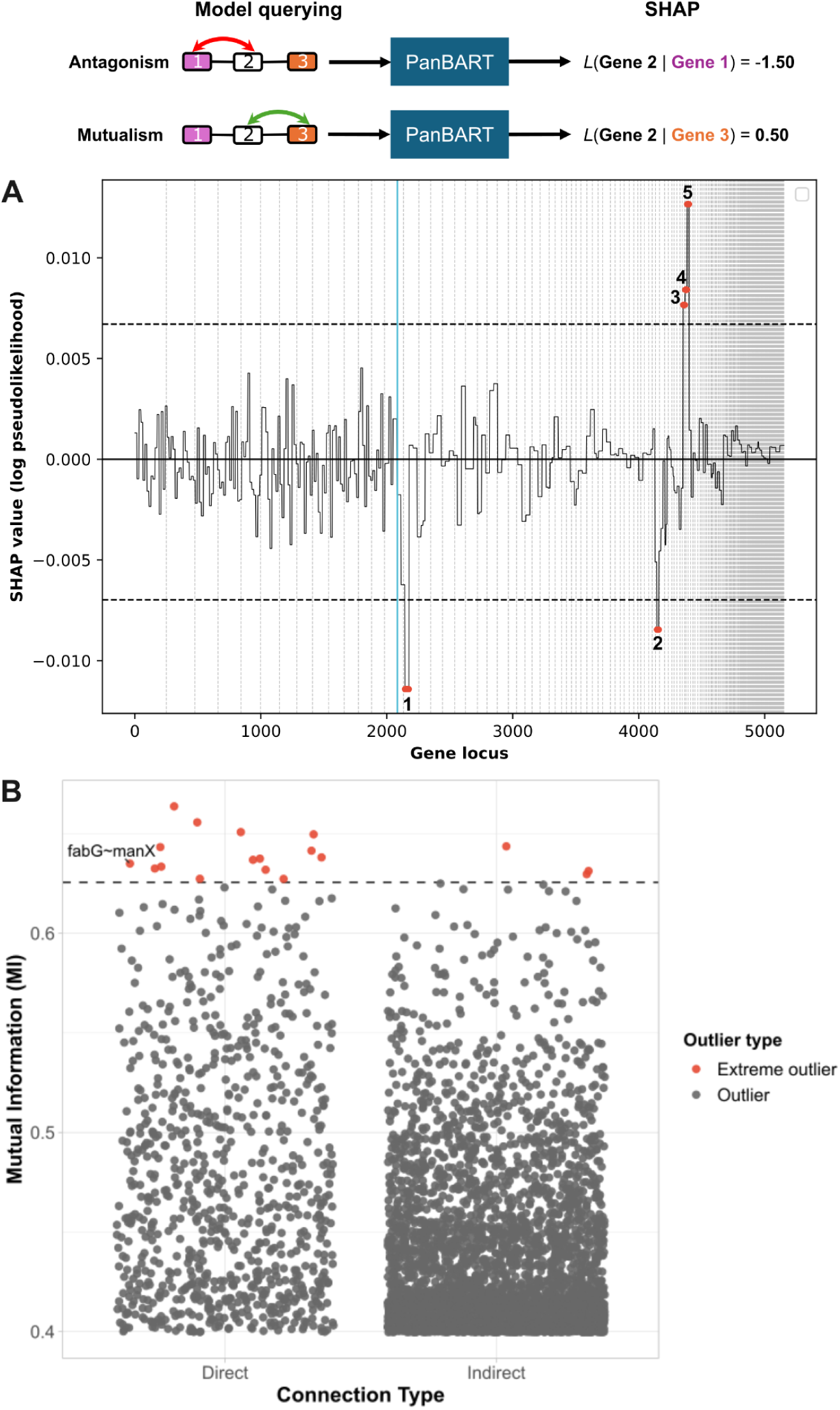
Co-selection detection in pangenomes. Genes under mutualistic and antagonistic co-selection should generate high and low SHAP values respectively (*top*). Gene-level co-selection detection (*bottom*). (**A**) SHAP (SHapley Additive exPlanations) values calculated from PanBART model of *E. coli* genome SAMEA4921639 from AllTheBacteria. The blue vertical line denotes the *cblB* gene locus. The black solid horizontal line denotes baseline SHAP value. The black dashed horizontal lines denote Tukey extreme inter quartile range (IQR) outlier values (see **Methods**). The red dots denote loci with extreme outlier SHAP values associated with the *clbB* gene locus, with numbered labels detailed with gene annotations in **Supplementary Table 5**. (**B**) Outlier gene pairings detected with Spydrpick. Only pairings with mutual information (MI) values above 1.5×IQR are shown, with pairings with MI values above threshold 3×IQR highlighted in red. Couplings are separated by whether they were identified as direct or indirect by ARACNE. Only gene pairings where both genes were annotated with gene names by Bakta are labelled.

## Discussion

Transformer models represent high-dimensional biological data in the form of genome or protein sequences in a lower dimensional feature space. This feature space reduction provides transformer models with unmatched generalisability to a multitude of bioinformatic tasks, such as functional annotation and phenotypic classification. However, applications of transformer models have not yet extended to genomic epidemiology tasks, such as differentiating lineages, identifying emergent lineages and determining the likelihood of outbreaks of drug-resistant lineages, requiring training on pathogen-specific data.

In this work, we trained a transformer model, PanBART, using gene content and gene order in species-specific bacterial pathogen datasets. We applied PanBART to five genomic epidemiology and evolutionary analysis tasks, benchmarking it against existing state-of-the-art methods on two bacterial pathogens with distinct population structures and different levels of pangenome diversity. PanBART training was fully unsupervised, meaning that it was not provided with any population-level information, such as inter-genome distances or phylogenetic relationships. Even without this information, PanBART accurately learned a representation of population structure based on gene content and gene order, clustering genomes belonging to the same lineages with higher agreement with PopPUNK than Sketchlib, which uses gene content only. This embedding space can also be used for lineage assignment of novel genomes, typically conducted during routine genomic surveillance or outbreak detection (Lees et al. 2019; Croucher and Didelot 2015). Genomic surveillance typically requires accurate and fast analysis; we show that PanBART lineage assignment is largely concordant with PopPUNK results, in comparable runtime to Sketchlib, whilst not being sensitive to dataset size. Furthermore, PanBART can be used to identify previously unseen lineages with high accuracy, and can generalise to classify closely-related species by leveraging sensitivity of transformer models to out-of-distribution data (Shi et al. 2025). PopPUNK can also identify unseen lineages which cluster separately to known lineages, although it is not designed for clustering of different species due to its sensitivity to large inter-genome distances. Therefore, PanBART generalises lineage clustering to closely-related species, unavailable in current state-of-the-art genomic epidemiology methods.

We investigated the effect of dataset sampling on lineages classification tasks with PanBART. We observed only minor performance impacts when training PanBART on 10% of the total training data compared with a full dataset. Therefore, small representative datasets of >10,000 genomes can viably be used to train species-specific PanBART models, which would include 18/24 of the WHO priority pathogens (**Supplementary Table 6**, excluding *Enterobacter, Citrobacter, Proteus, Serratia* and *Morganella* species based on respective pathogen AllTheBacteria assemblies counts (Sati et al. 2025; Hunt et al. 2024)). Future work will expand the number of trained PanBART models to include these high-risk pathogens.

We further explored the epidemiological applications of PanBART that go beyond existing genomic epidemiological analysis tools. We leveraged the higher-order gene-gene interactions captured by PanBART to enable accurate prediction of gene uptake potential in *E. coli*, akin to *in silico* mutational screens previously explored with transformers trained on protein sequences (Meier et al. 2021; Brandes et al. 2023). This feature is not available in existing genome-based machine learning approaches, as they rely on a few informative features at inference time (Biffignandi et al. 2024; Lees et al. 2020; Beavan et al. 2024). While lineage classification, and therefore population structure, had a clear effect on distinguishing lineages with high and low risk of gaining *blaCTX-M*, it did not fully explain the capability of PanBART to classify lineage risk. Genomes from ST73 and ST95 naturally containing *blaCTX-M* were classified as high-risk, despite belonging to low-risk lineages. Therefore, PanBART identified additional genome features predisposing these genomes to *blaCTX-M* uptake, which were not shared with other members of these low-risk lineages. This result provides the groundwork for further *in silico* and empirical analysis to ascertain the identity of the genes associated with *blaCTX-M* uptake, and identify the presence of higher-order gene-gene interaction networks that support acquisition of drug resistance. Analysis of gene uptake potential can also be easily extended to other clinically-relevant genes, such as those involved in virulence, vaccine-escape or other antibiotic resistance mechanisms.

We additionally explored detection of gene-level co-selection, which has been achieved previously using non-tranformer methods (Whelan et al. 2020; Beavan et al. 2023). We were able to identify a previously unseen, but theorised, association between a siderophore and an iron-regulated bacteriocin (Martin et al. 2013; Wallenstein et al. 2020). We therefore believe that PanBART enables sensitive detection of a weak signal co-selection, as it is run on the level of individual genomes, rather than on whole populations, where the signal may be diluted by population structure. However, as PanBART was only able to identify this relationship in a single genome and therefore itself is likely sensitive to noise from population structure and assembly quality, we believe co-selection detection using PanBART should be strictly used as an exploratory tool, with experimental validation being necessary to verify detected relationships.

We observed notable differences in the embedding space of *S. pneumoniae* and *E. coli*, and in turn a difference in performance of lineage clustering and assignment between the two species. Population structure is a consequence of the evolutionary history of a population, including selective processes, horizontal gene transfer and demography (Harrow et al. 2021; Rocha 2018; Baumdicker and Kupczok 2023). As PanBART is not provided with any phylogenetic information, its ability to represent the populations of two pathogens with distinct population structures and levels of pangenome diversity indicates that information on population structure, and therefore an abstraction of the evolutionary processes giving rise to population structure, is learned by the model. Therefore, transformer models may be useful in understanding how gene-gene interactions have driven the formation of pathogen populations we observe today (McInerney 2026; Neher and Shraiman 2009; Cummins et al. 2026). Combination of gene pseudolikehoods provided by transformer models with phylogenetic reconstruction approaches may provide insight into how higher-order gene-gene interactions have shaped past and extant bacterial populations.

An obvious limitation of this approach is the computational efficiency of model training, requiring several high-performance GPUs for multiple days. However, model training is required only once per species, with inference tasks, such as lineage assignment, then running rapidly at rates of hundreds of genomes per minute. Additionally, token dictionaries were necessarily limited to genes over 0.1% frequency, meaning that rare genes below this frequency were not considered. Recent work has shown that in some cases rare genes are maintained by selection (Douglas and Shapiro 2024), meaning that these genes may hold biologically relevant information that could be learned by the model to further improve task accuracy. Additionally, our interpretation of token functionality is limited to gene-level rather than nucleotide-level, meaning that PanBART cannot be used on species with limited structural or gene-level variation, such as *Mycobacterium tuberculosis (Behruznia et al. 2025)*. This design choice was made as transformer models are limited to a context length of thousands of tokens due to attention matrices scaling quadratically with content window length. While we applied a long-context transformer to mitigate this issue (Beltagy et al. 2020), our work could be further extended to combine sequences within genes as well as their order within more recent models with even longer context windows, such as state-space models (Nguyen et al. 2024, 2023) or long-term memory recurrent neural networks (Behrouz et al. 2025, 2024).

The models trained here were not explicitly corrected for population structure, which is observed due to strong linkage between variants in asexually reproducing species (Lees et al. 2016; Whelan et al. 2020; Pensar et al. 2019). We found evidence that PanBART does not exclusively learn population structure when applied to epidemiological tasks, exemplified by high-risk ST73 and ST95 genomes being correctly identified for *blaCTX-M* uptake, despite belonging to low-risk lineages. However, the lack of expected associations in co-selection analysis may be due to a high phylogenetic signal masking the signal of gene-gene co-selection (Pensar et al. 2019). There is currently no standardised means of accounting for population structure when training genomic transformer models, and further work could focus on exploring ways of distinguishing phylogenetic and co-selection signals between loci, such as with sequence reweighting used in protein contact map detection and allelic co-occurance detection (Morcos et al. 2011; Pensar et al. 2019).

Transformer models provide highly flexible frameworks for genome analysis, greatly extending the analysis capabilities provided by conventional machine learning approaches. We show a bidirectional long-context model, PanBART, is a versatile tool for epidemiological analysis. We detail applications of PanBART to a range of epidemiological analyses, where it performs equivalently or better than existing tools, and provides a wide breadth of functionality in a single model. Our work lays a foundation for further use of transformer models in epidemiological analysis of bacterial pathogens, which will greatly benefit from the growing availability of universally-assembled and centrally-stored genome dataset repositories.

## Materials and Methods

### Dataset collation and stratification

Genomes for model training and analysis were gathered from AllTheBacteria (Hunt et al. 2024) (release 0.2, increment 2024-08). Genomes annotated as “*Escherichia coli*”, *“Streptococcus pneumoniae*” and *“Streptococcus mitis*”, as well as having “high quality” labels were extracted from metadata provided with AllTheBacteria genomes (≥99% reads have same single species assignment, ≥90% completeness and ≤5% contamination from CheckM2 (Chklovski et al. 2023), total assembly length between 100kbp and 15Mbp, maximum number of contigs 2,000, minimum N50 2,000).

Genomes for *S. pneumoniae* and *E. coli* were then assigned to sequence clusters, denoting lineages, using PopPUNK v2.7.6. For *S. pneumoniae*, genomes were sketched (‘poppunk ––create-db ––length-sigma 1’) and run through the PopPUNK quality control workflow (‘poppunk ––qc-db ––max-pi-dist 0.03 ––max-a-dist 0.4’), to which an existing cluster assignment model (*S. pneumoniae* v10, available from https://gps-project.cog.sanger.ac.uk/GPS_v10.tar.gz) was fitted (‘poppunk ––use-model’). For *E. coli*, an iterative cluster assignment process was used; the dataset was split into eight batches, to which an existing cluster assignment model (*E. coli* v2, available from https://ftp.ebi.ac.uk/pub/databases/pp_dbs/escherichia_coli_v2_refs.tar.bz2) was fitted (‘poppunk_assign ––run-qc ––run-qc ––max-zero-dist 1 ––max-merge 3 ––update-db fast ––overwritè). This generated a cluster assignment file for each batch, which were then merged into a single cluster assignment file using a custom script (pangenome_LLM/merge_iter_poppunk.py).

Datasets were additionally subset and stratified to emulate model training on incompletely sampled populations. For both species, the number of each cluster was counted, and 10 clusters were removed completely for training of models with a subset of the total lineages, with sizes spanning each 10% quantile (see **Supplementary File 1**). Datasets were also sampled to 10% of the full dataset size using the script pangenome_LLM/sample_PopPUNK.py. This resulted in 4 datasets per species, detailed in **Supplementary Table 2**.

For each dataset, genomes were tokenised using WTBCluster v0.1.0 (https://github.com/samhorsfield96/WTBcluster), a Snakemake pipeline (Köster and Rahmann 2012) we wrote for generating integer-representations of bacterial genomes. Using WTBCluster, genomes were annotated using Pyrodigal v3.6.3 (Larralde 2022) to predict coding sequences, before being clustered using MMSeqs2 Linclust v16.747c6 (Steinegger and Söding 2017, 2018). Each gene cluster was assigned an integer, and each genome converted to a string of integers representing the ordering of each gene cluster in the genome. Contig breaks were represented as ‘_’, and genes on forward or reverse strands were represented as positive or negative integers respectively. Clustering was performed at 50% identity and 50% coverage (coverage mode [‘--cov-modè]: 0, cluster mode [‘--cluster-modè]: 2, identity mode [‘--seq-id-modè]: 2, alignment mode [‘--alignment-modè]: 3). For *S. pneumoniae*, all coding sequences were clustered in a single step for each species (‘mmseqs2_num_batches’: 1), whilst for *E*. *coli*, this was done in two steps (‘mmseqs2_num_batches’: 2). For each dataset, rare tokens under 0.1% frequency were removed using a custom script (pangenome_LLM/remove_rare_tokens.py) to reduce the token corpus size required during model training and inference.

For each species, tokenised genomes were split into three groups: training, used for PanBART training; validation, used during training to calculate per-epoch loss and accuracy and guide early stopping of training; and testing, held out completely during training and used for downstream analysis. Datasets were stratified into training, validation and testing based on PopPUNK cluster assignment labels, with each cluster being split into 80%:10%:10% splits for training, validation and testing respectively using a custom script (pangenome_LLM/stratify_data.py)

### PanBART training

PanBART uses an existing model architecture, the Longformer Encoder-Decoder (Beltagy et al. 2020), which allows for greater computational efficiency versus conventional transformer architectures, through implementation of sliding-window attention in contrast to full-window attention. A full attention window, used in conventional transformers, captures all token-token interactions across the full window, in this case all gene-gene interactions across the full genome. This is computationally expensive, as all pairwise interactions must be stored in GPU memory, and so only scales to ∼2,000 tokens in a single context window. In contrast, a sliding-window attention, used in Longformer, captures gene-gene interactions in smaller slices across the full genome, and so scales to larger windows of >10,000 tokens in a single context window. We used Transformers v4.42.3 (Wolf et al. 2019) to access the Longformer Encoder-Decoder architecture.

Stratified tokenised genome datasets were provided directly to PanBART as training and validation data, with PanBART being trained separately for each species and dataset sample. To prevent PanBART overfitting to arbitrary contig ordering, contig order and orientation was randomised for each genome in each epoch. If contig order were not randomised, the model would implicitly learn how contigs are arranged, meaning it would not generalise to assemblies with another arbitrary contig order.

For both *S. pneumoniae* and *E. coli* datasets, PanBART was trained with a sliding window length of 512 tokens, batch size of one, eight attention heads, eight attention layers per head, a hidden embedding layer size of 256 (‘--attention_window 512 ––batch_size 1 ––num_heads 8 ––num_layers 8 ––embed_dim 256’). The learning rate was set to 1×10^-5^ (‘--learning_rate 1e^-5^’), to which a L2 normalisation was applied of 1×10^-4^ to prevent overfitting (‘--weight_decay 1e^-4^’). We found training with a batch size of one produced good results (**Supplementary Figure 2**), while also keeping computation requirements to a minimum, in line with previous work (Marek et al. 2025). During encoder training, 15% of tokens were randomly masked from each genome (‘--prop_masked 0.15’) in line with previous work (Devlin et al. 2018; Lewis et al. 2019). During each epoch, 40% of nodes in hidden layers were dropped to reduce the effect of model overfitting to large closely-related lineages (‘--model_dropout_rate 0.4’), which would otherwise result in the model performing well on common lineages, and poorly on rare lineages. Early stopping was enabled, which ended training when the loss did not change by a proportion of 0.01 for 10 consecutive epochs (‘--early_stop_patience 10 ––min_delta 0.01’). PanBART was trained with a maximum context window length of 8192 tokens (--max_seq_length 8192) and 9728 (‘--max_seq_length 9728’) for *S. pneumoniae* and *E. coli* respectively. This was implemented in the script panBART/panBART.py.

### Embedding visualisation

PanBART embeddings were generated by passing genomes through trained PanBART models, gathering final layer embedding vectors for each token generating a mean pooling of vectors (panBART/compute_sequence_embedding.py). Embeddings were visualised using UMAP (pangenome_LLM/plot_embeddings.py), with the top 10 largest PopPUNK clusters coloured.

Sketchlib embeddings were generated using pp-Sketchlib v2.1.4 (Lees et al. 2022). Sketches were generated for training and testing datasets using a sketch size of 10000 and k-mer sizes ranging from 13bp to 29bp with an interval of 4bp (‘Sketchlib sketch –s 10000 –k 13,29,4’). Sparse pairwise distance matrices were generated from sketches with 1000 nearest neighbours per genome (‘Sketchlib query sparse infile ––kNN 1000 ––accessory’). Sparse pairwise distances matrices were then converted to embeddings and plotted as UMAP plots using a custom script (pangenome_LLM/plot_embeddings_sparse.py). The top 10 largest PopPUNK clusters were coloured as before.

PanBART and Sketchlib embeddings were quantitatively compared using Silhouette scores implemented in Scitit-learn v1.6.1 (Pedregosa et al. 2011). Silhouette scores describe the agreement between inter– and intra-cluster distances based on an external labelling scheme. PopPUNK cluster labels were used as external labels and applied to PanBART and Sketchlib embeddings.

### Lineage assignment

PanBART and Sketchlib embeddings from training data were used to assign PopPUNK cluster labels to testing data. k-nearest neighbours (kNN) classifiers were trained for each species and dataset, with known PopPUNK cluster labels for testing data being compared to predicted labels to calculate the accuracy of cluster assignment (panBART/assign_cluster.py for PanBART, pangenome_LLM/assign_strains_sparse.py for Sketchlib). Balanced accuracy was calculated using “balanced_accuracy_score” in Scitit-learn, and was used to compare performance of PanBART and Sketchlib embeddings. Balanced accuracy is defined as the average recall obtained for each class and therefore removes potential bias caused by imbalanced class sizes.

### Whole genome pseudolikelihood analysis

To generate whole genome pseudolikelihoods, query genomes were passed through trained PanBART models using both the encoder and decoder. We randomly sampled 50 query genomes from lineages included in and held out of *S. pneumoniae* training data (see **Supplementary File 1**). The logit of each gene in the sequence was extracted from the final classification layer of the decoder. This layer is a probability distribution over all possible token values for a given position in a genome, and is used by the model to predict the identity of a masked token by taking a weighted sample of N=1 using this distribution. As we knew the gene identity for a given query genome, acquiring the respective logits for each gene token enables measurement of model confidence for a given token at a given position. Known gene logits from all loci for each query genome were summed to generate a final whole genome pseudolikelihood. The analysis was conducted using the script panBART/pseudolikelihood.py. We used log pseudolikelihoods for genomes used in training as the initial benchmark, as we expected these to be highest due to the model having been trained on this dataset.

### Token conditional pseudolikelihood analysis

To generate gene token pseudolikelihoods conditioned on the genomic background, target tokens (e.g. tokens functionally annotated as *blaCTX-M*) were first removed from each genome if present. The target token was then iteratively added at each position and passed through a trained PanBART model. The logit of the target token at that given position was then extracted, producing a logit value for the target token being found at every location in the genome. The position with the highest logit value was then taken as the most likely insertion sequence of the target token (panBART/compute_token_likelihood.py).

For the analysis of *E. coli* association with *blaCTX-M*, genomes from the NORM project (Gladstone et al. 2021) were assigned to PopPUNK clusters as described above. Sequence types were associated with PopPUNK clusters using these genomes, enabling identification of genomes in the AllTheBacteria that belonged to ST131 B/C, ST131 A, ST95 and ST73. Genes in each genome were annotated using Bakta v1.9.4 (database v5.1) (Schwengers et al. 2021) and assigned to tokens (pangenome_LLM/annotate_tokens.py), with gene annotations being used to identify tokens annotated as *blaCTX-M*. Genomes from each sequence type were assigned as either *blaCTX-M* positive (+ve) or negative (-ve) based on the presence of the token identified as *blaCTX-M*. These stratified datasets were then used in token conditional pseudolikelihood analysis. Ten genomes from each stratified dataset were used in analysis.

### Co-selection analysis

SHAP (SHapley Additive exPlanations) analysis was used to identify evidence of co-selection between genes in individual genomes. *E. coli* genomes were first selected from different PopPUNK clusters using a custom script (pangenome_LLM/stratify_N_genomes.py), with two genomes per cluster being selected which were annotated as containing the *cblB* gene. SHAP values were then generated for each genome using the SHAP python package v0.48.0 (Lundberg and Lee 2017). We used the partition explainer method on the encoder only, using non-target tokens as features and the logit of the target token as the dependent variable. Briefly, the partition explainer method augments the non-target tokens by iteratively masking portions of the genome, and determines the impact on the logit of the target token. Masking regions that have a large impact on the target token logit indicates that these regions are important to the model when predicting the target token at that position, and can be positive in cases where a non-target and target tokens are predicted to be be found together, and negative in cases where they are predicted to be mutually exclusive. Genes were identified as outliers based on Tukey extreme outlier values (Hoaglin et al. 2000), *T* = *Q*_3_ + (3 × (*Q*_3_ – *Q*_1_), where *T* is the extreme outlier threshold, *Q*_1_ and *Q*_3_ are the lower and upper quartiles respectively. To prevent genes from being associated due to linkage, a 65 gene distance cut off was used, with gene pairs below this distance from the target not being identified as outliers (based on (Mallawaarachchi et al. 2024), linkage in *E. coli* ∼65 kb assuming genes are ∼1 kbp in length) (panBART/compute_SHAP.py for SHAP calculation, pangenome_LLM/plot_SHAP_publication.py for plotting).

Co-selection was also analysed using Spydrpick v1.2.0 (Pensar et al. 2019). Tokenised *E. coli* genomes used in PanBART training were first converted into a gene presence/absence matrix using a custom script (pangenome_LLM/generate_pa_matrix.py). This presence/absence matrix was then converted to a multiple-sequence alignment, with “c” denoting a gene being present, and “a” denoted a gene being absent, as used previously for co-selection detection using unitigs (Kuronen et al. 2024), using the script pangenome_LLM/generate_MSA_from_pa_matrix.py. Spydrpick was then run on these gene-level alignments using a minimum allele frequency of 0.05 and a maximum of 100 million mutual information (MI) values (pangenome_LLM/analyse_spyderpick.py), returning direct and indirect interactions identified by ARACNE with no minimum MI values. Only gene pairs which both had non-hypothetical annotations from Bakta v1.9.4 (Schwengers et al. 2021) were annotated.

## Code availability

Code for PanBART v0.1.0 used for training and downstream analysis of model outputs is available on Github (https://github.com/samhorsfield96/panBART) and Zenodo (doi: 10.5281/zenodo.19814981). Code for downstream analysis not directly associated with PanBART models, collectively named “pangenome_LLM”, is also available on Github (https://github.com/samhorsfield96/pangenome_LLM) and Zenodo (doi: 10.5281/zenodo.19815224). All models and tokenizers are available on Zenodo (doi: 10.5281/zenodo.19823911).

## Acknowledgements

We would like to thank Dr. Tommi Mäklin at the University of Oslo for his help identifying the *clb* locus as a useful test-case for co-selection detection in *E. coli*.

## Author Contributions

Conceptualization: S.T.H., S.D.B, C.C. and J.A.L. Methodology: S.T.H. Software: S.T.H. and J.O.M. Validation: S.T.H. Formal analysis: S.T.H. Investigation: S.T.H. Resources: J.A.L. Data curation: S.T.H. Writing – original draft: S.T.H. Writing – review and editing: All authors. Visualization: S.T.H. Supervision: J.A.L. Project administration: S.T.H. and J.A.L. Funding acquisition: S.D.B, C.C. and J.A.L.

