## Supplementary Material for "Species-specific transformer models of bacterial gene order and content for genomic surveillance tasks"

#### Supplementary Figures

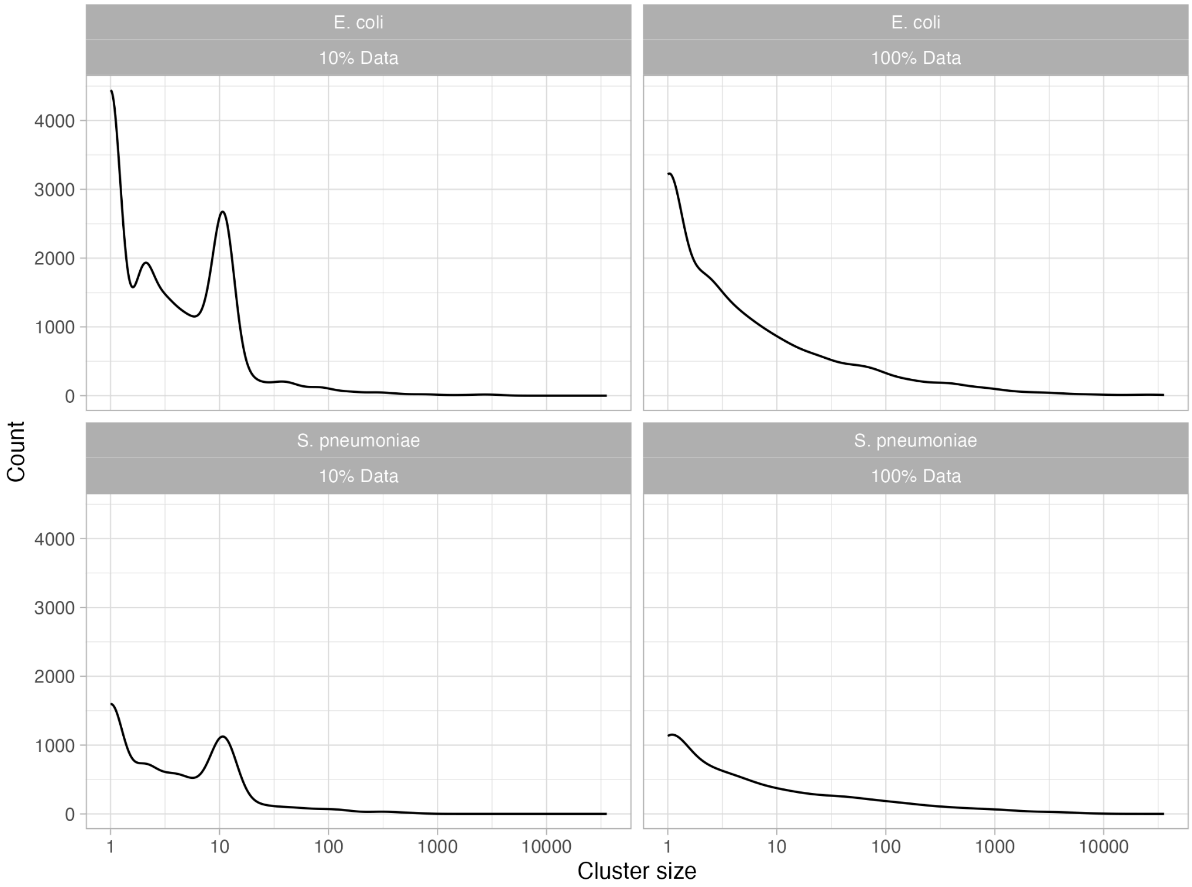

Supplementary Figure 1: PopPUNK cluster size density plot for all genomes used in analysis. Data shown for 10% sample and full (100%) datasets.

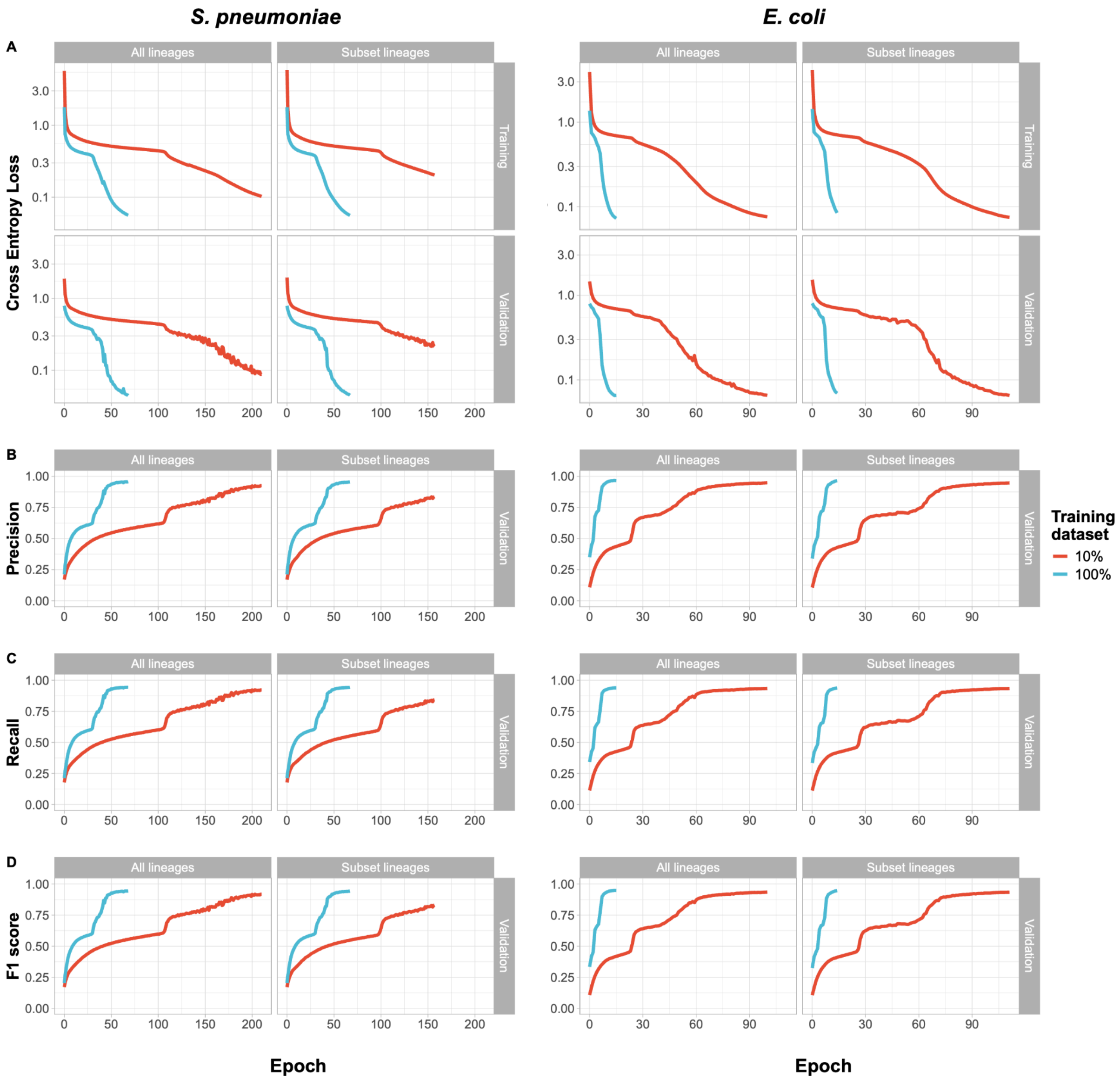

Supplementary Figure 2: PanBART accuracy during model training for *S. pneumoniae* (left) and *E. coli* (right) datasets. Four models were trained for each species with different subsets of data. Datasets were split based on the number of PopPUNK lineage clusters they contained: all lineages, all PopPUNK lineage clusters were included in training; subset lineages, a selection of PopPUNK lineage clusters, with a distribution of sizes, were held out of training. Datasets were also split by the proportion of the total dataset they contained, either 100% (blue lines) or 10% (red lines) of genomes. (**A**) Cross entropy loss per epoch for training and validation datasets. (**B**) Precision of token predictions at each masked position. (**C**) Recall of token predictions at each masked location. (**D**) F1 score of token predictions at each masked location.

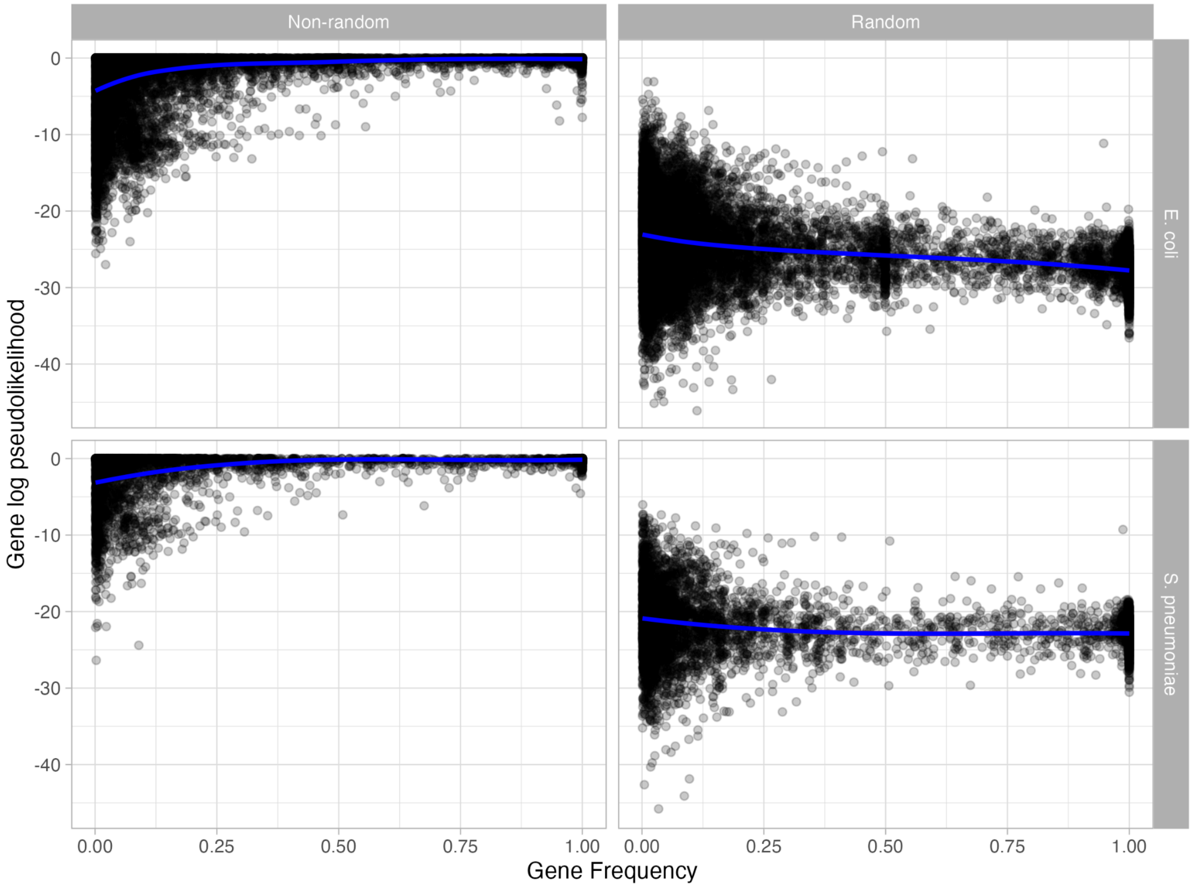

Supplementary Figure 3: Effect of randomising gene order on relationship between gene frequency and gene log pseudolikelihood. Data is shown for 10 randomly selected genomes from the training datasets for models trained on 100% of available genomes using all lineages from each species. The same 10 genomes were used across non-randomised and randomised analysis. The blue trendline and red 95% confidence intervals were generated using the LOESS regression.

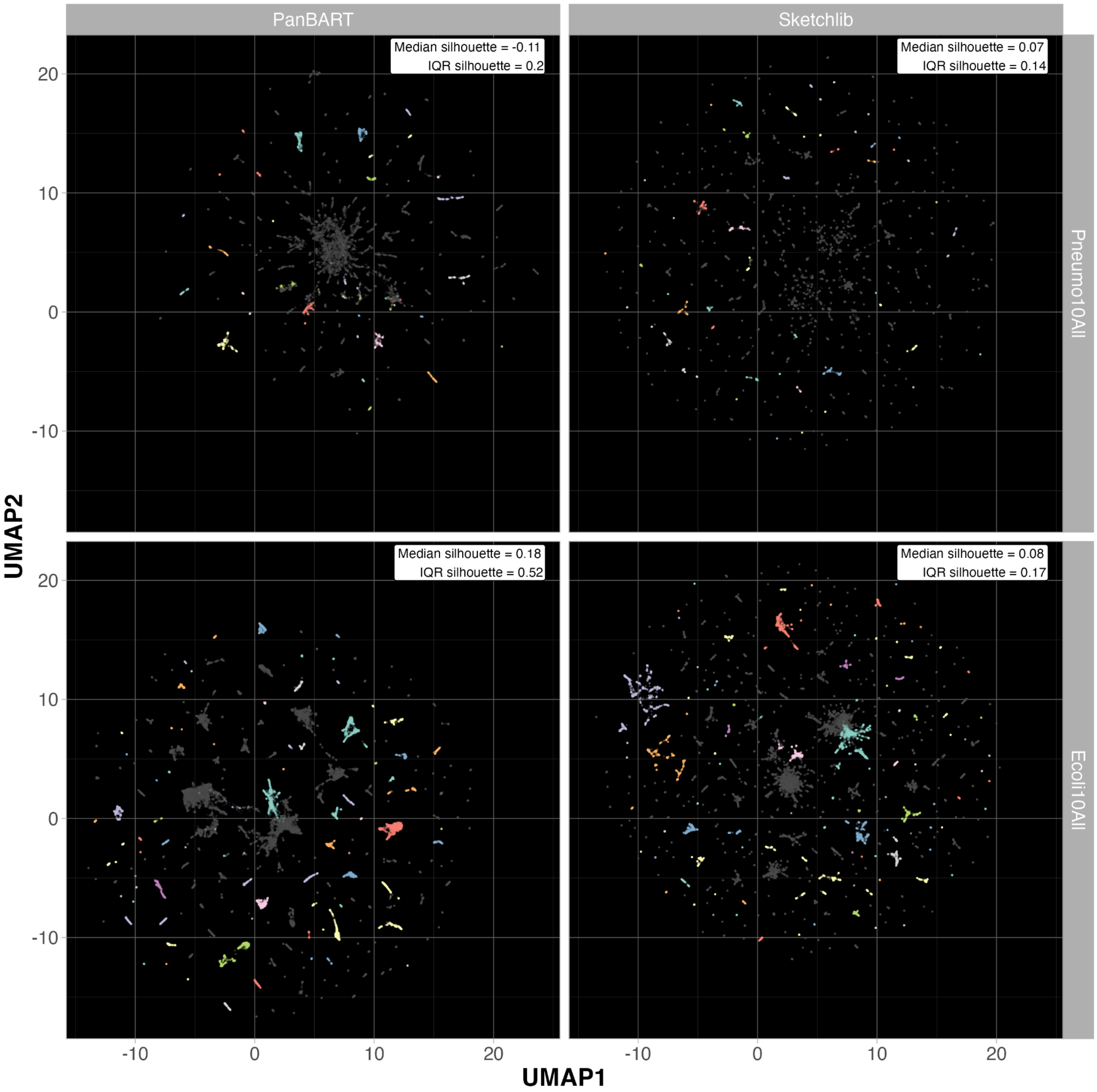

Supplementary Figure 4: Comparison of PanBART (left) and Sketchlib (right) genome embeddings. Each point represents the embedding of a single genome. *S. pneumoniae* (top) and *E. coli* (bottom) for the models trained on all PopPUNK lineage clusters, with 10% of the total training data. Embeddings are shown for genomes used in model training. Colours show the 10 largest PopPUNK lineage clusters for each species. The median and Inter-Quartile Range (IQR) of silhouette scores for the 10 largest PopPUNK lineage clusters is shown in the top right corner of each plot.

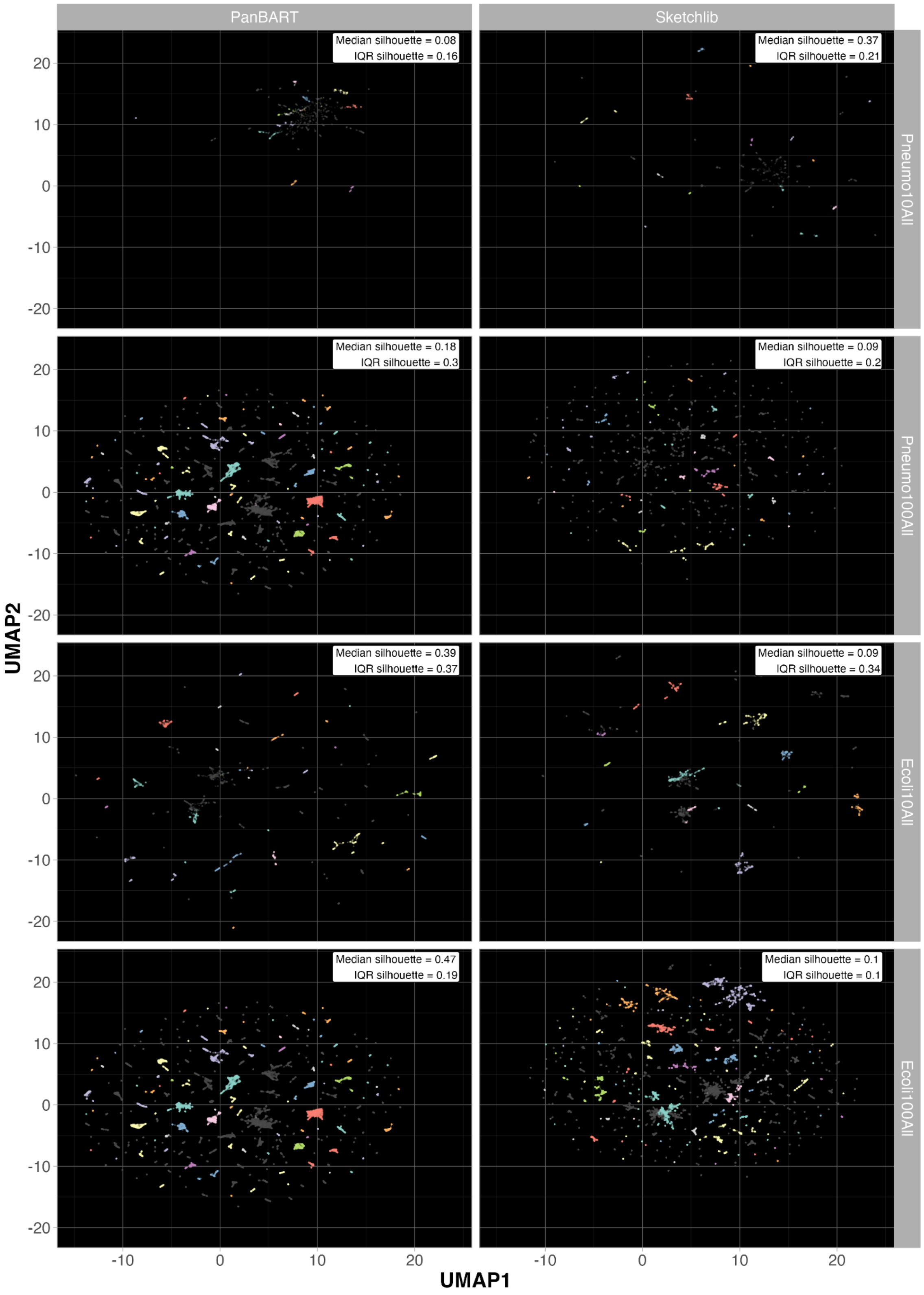

Supplementary Figure 5: Comparison of PanBART and Sketchlib genome embeddings for testing data, where all lineages were used in model training. Each point represents the embedding of a single genome. Colours show the 10 largest PopPUNK lineage clusters for each species. The median and Inter-Quartile Range (IQR) of silhouette scores for the 10 largest PopPUNK lineage clusters is shown in the top right corner of each plot.

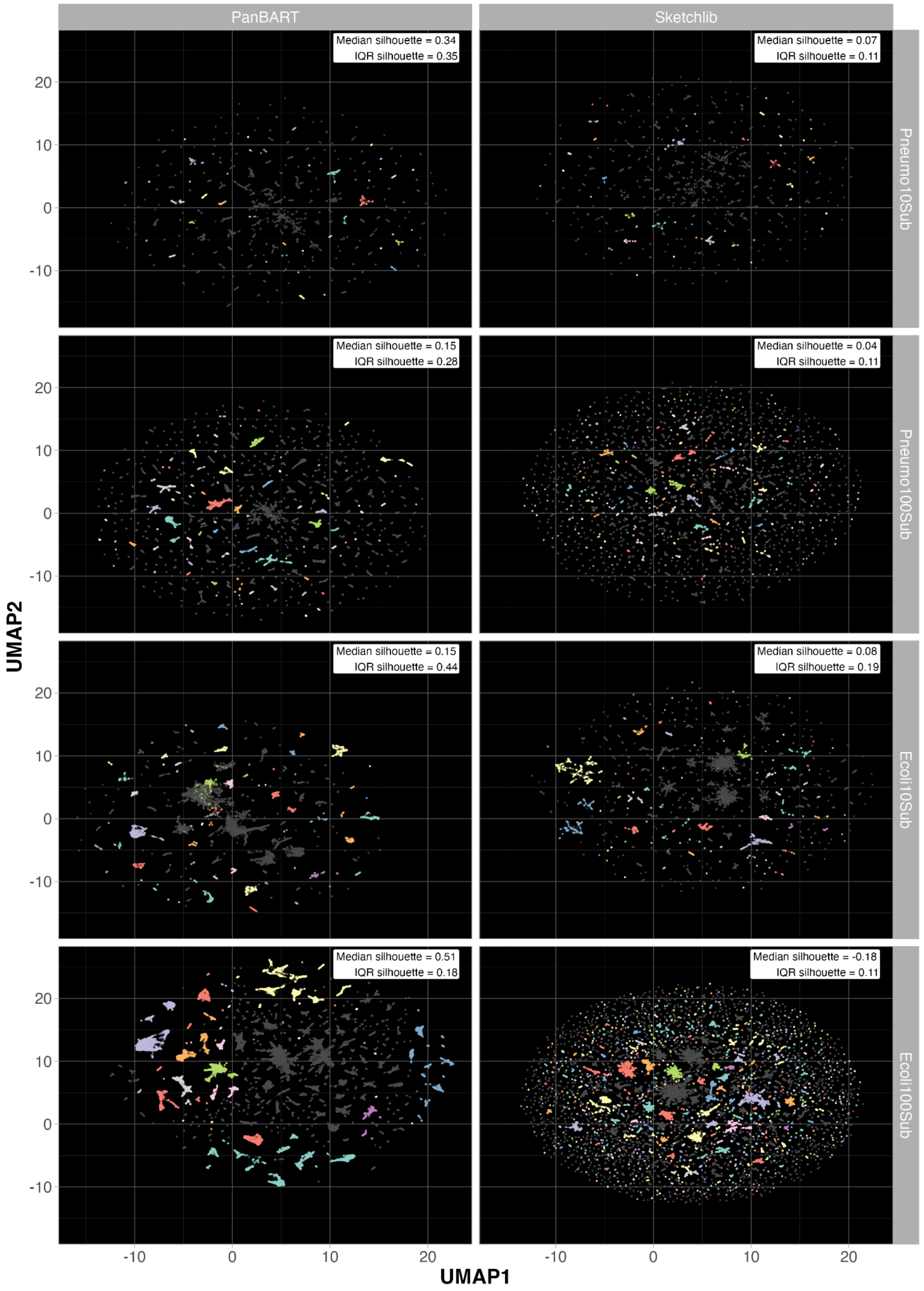

Supplementary Figure 6: Comparison of PanBART and Sketchlib genome embeddings for training data, where the model was trained on a subset of lineages. Each point represents the embedding of a single genome. Colours show the 10 largest PopPUNK lineage clusters for each species. The median and Inter-Quartile Range (IQR) of silhouette scores for the 10 largest PopPUNK lineage clusters is shown in the top right corner of each plot.

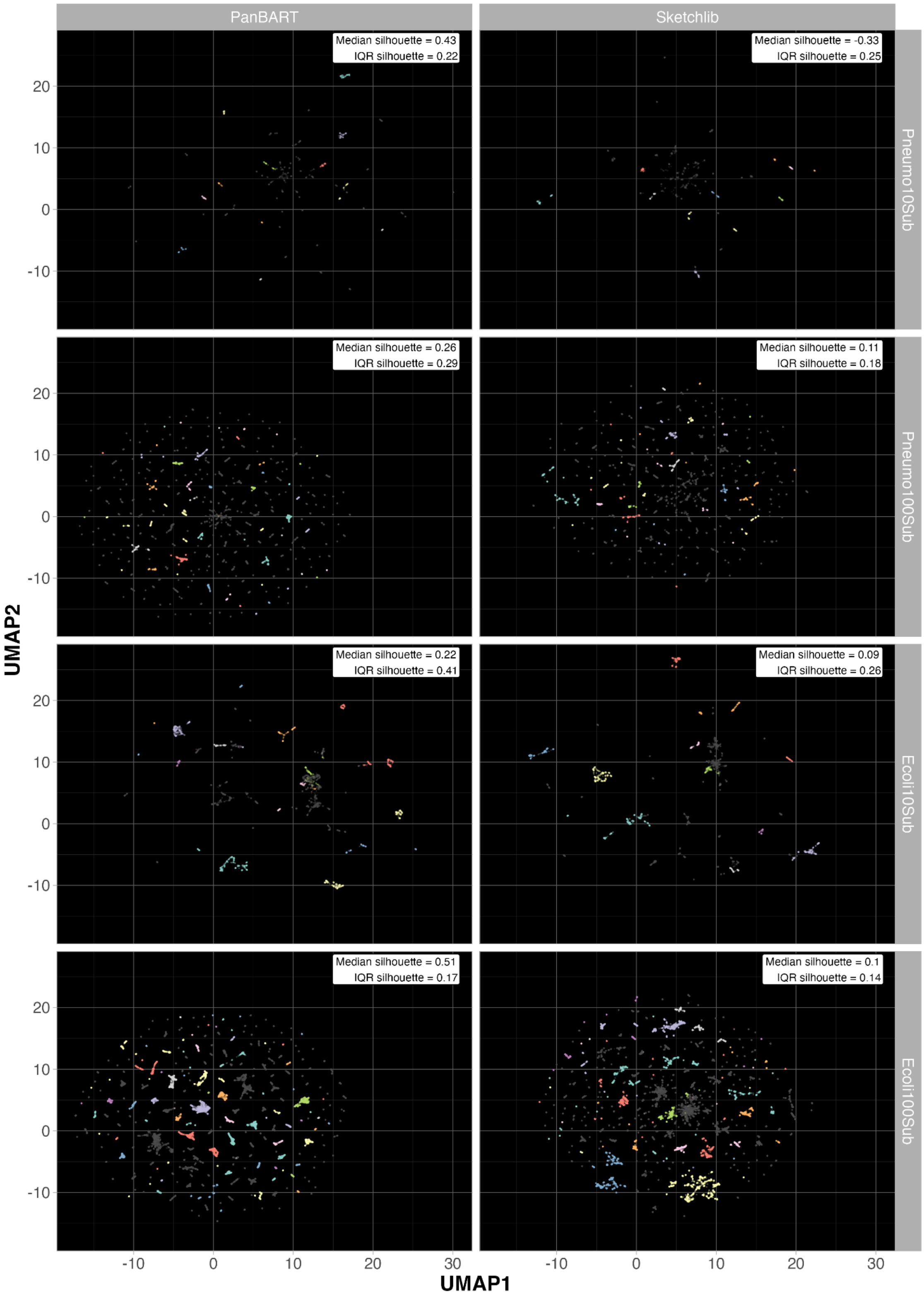

Supplementary Figure 7: Comparison of PanBART and Sketchlib genome embeddings for testing data, where the model was trained on a subset of lineages. Each point represents the embedding of a single genome. Colours show the 10 largest PopPUNK lineage clusters for each species. The median and Inter-Quartile Range (IQR) of silhouette scores for the 10 largest PopPUNK lineage clusters is shown in the top right corner of each plot.

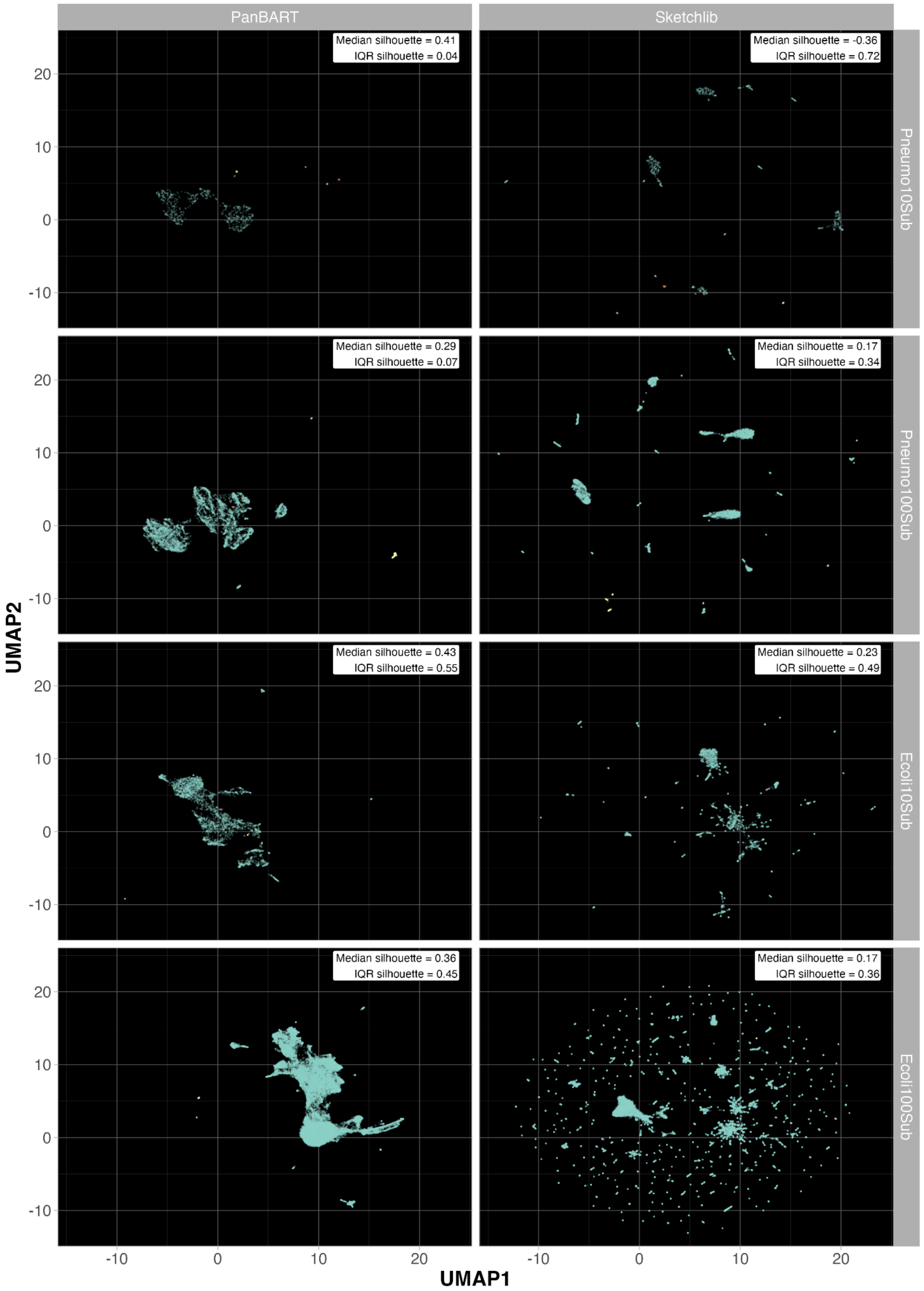

Supplementary Figure 8: Comparison of PanBART and Sketchlib genome embeddings for held-out lineages data, where the model was trained on a subset of lineages. Each point represents the embedding of a single genome. Colours show the 10 largest PopPUNK lineage clusters for each species. The median and Inter-Quartile Range (IQR) of silhouette scores for the 10 largest PopPUNK lineage clusters is shown in the top right corner of each plot.

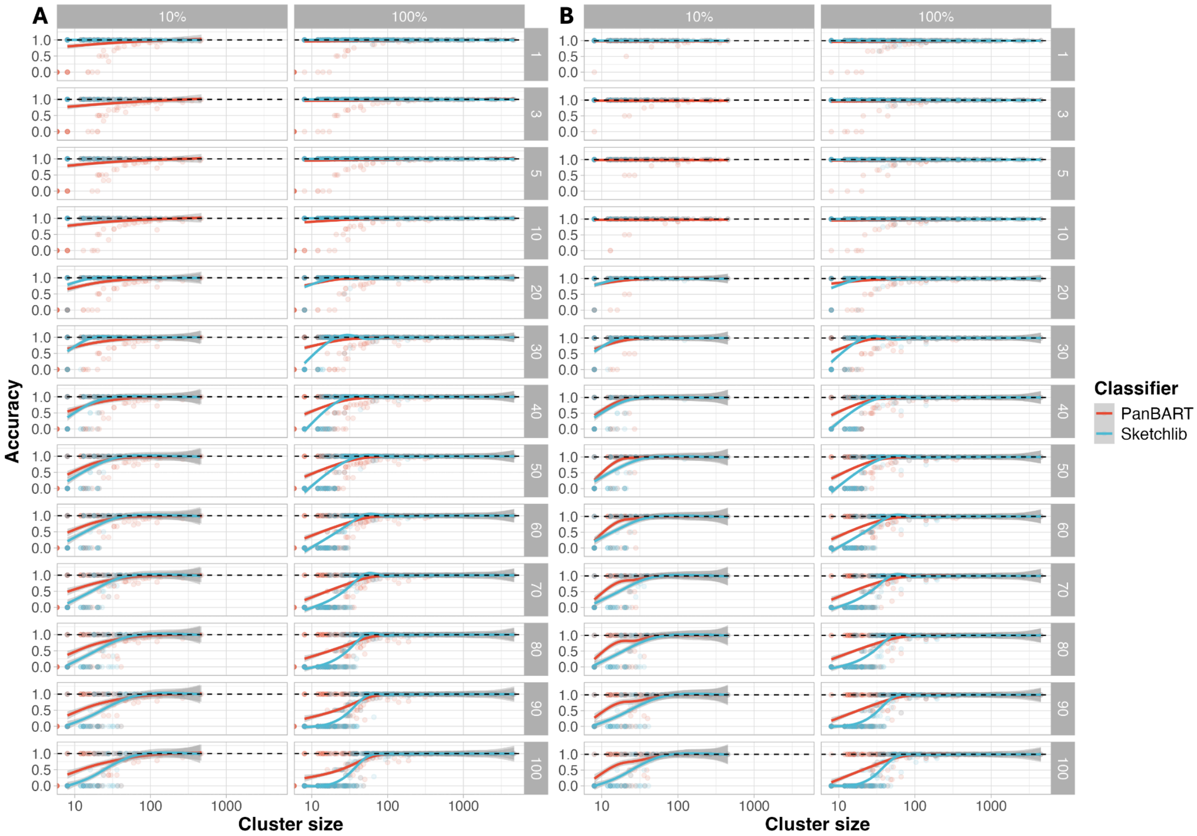

Supplementary Figure 9: *S. pneumoniae* accuracy of lineage assignment versus lineage cluster size for testing genomes for PanBART and Sketchlib for subsets of lineages (**A**) and all data (**B**). Horizontal facets describe training dataset size, vertical facets describe k used for kNN classification. Trend lines and error ranges were generated using a generalised additive model (GAM) with default parameters.

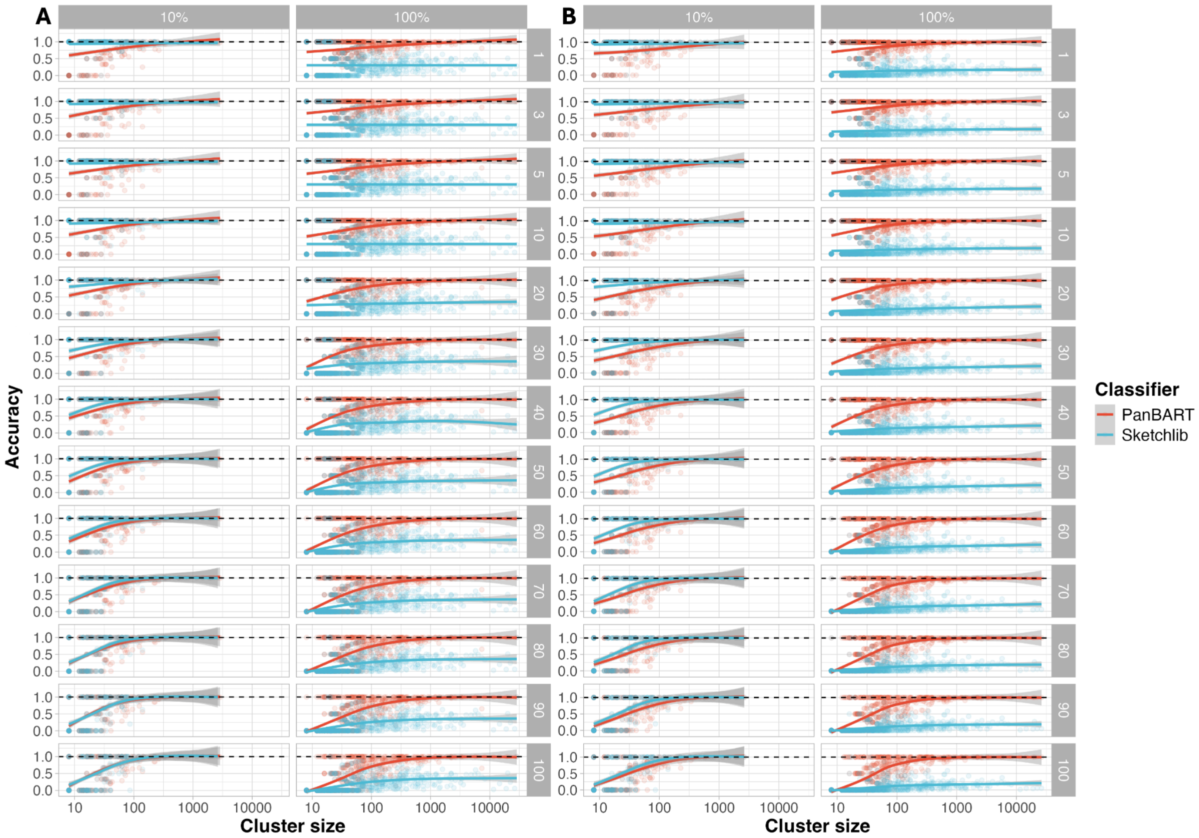

Supplementary Figure 10: *E. coli* accuracy of cluster assignment versus cluster size for testing genomes for PanBART and Sketchlib for subsets of lineages (**A**) and all data (**B**). Horizontal facets describe training dataset size, vertical facets describe k used for kNN classification. Trend lines and error ranges were generated using a generalised additive model (GAM) with default parameters.

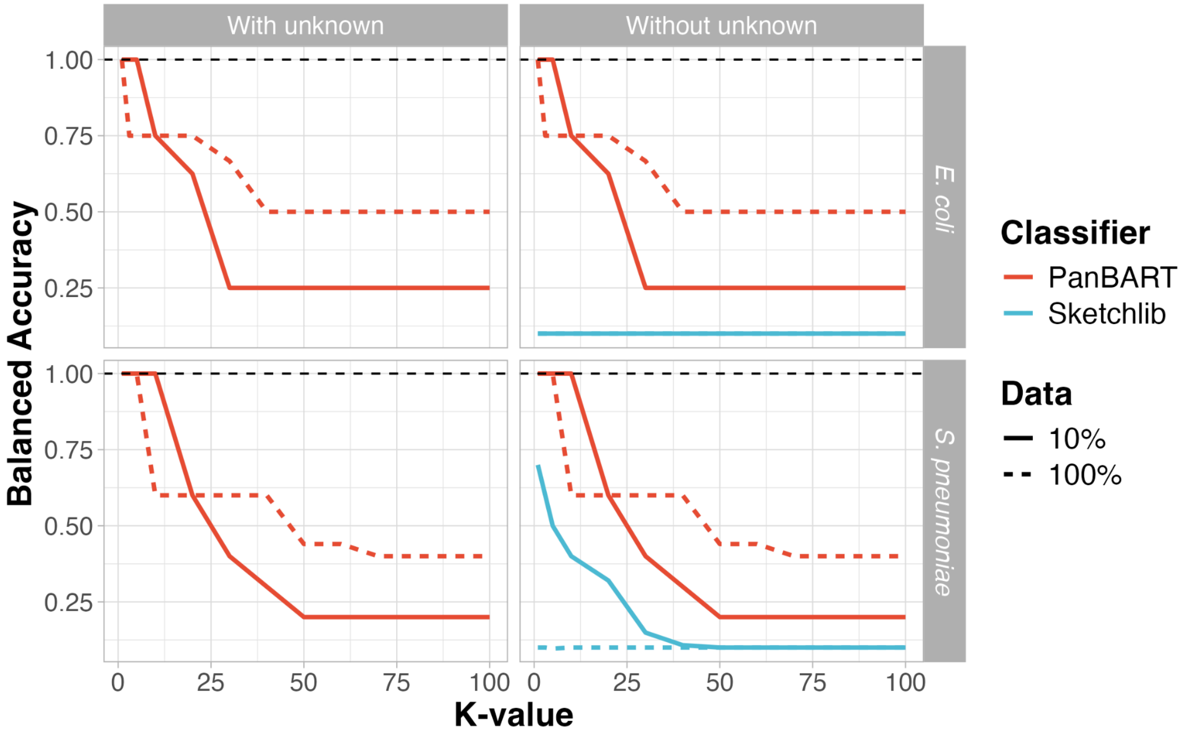

Supplementary Figure 11: Balanced accuracy comparison of lineage assignment of lineages held out of model training. Each line represents the balanced accuracy of PopPUNK lineage cluster assignment based on k-nearest neighbours (kNNs) in embedding space. PanBART results are shown by red lines; Sketchlib by blue lines. K-values used for classification are shown on the X-axis. Models were compared based on the dataset used for training: model training with either 10% (solid lines) or 100% (dashed lines) of available training data. Horizontal facets describe whether unknown tokens were attended to at inference time ("With unknown") or ignored from attention ("Without unknown"). Balanced accuracy for *E. coli* with Sketchlib was identical for both 10% and 100% of training data.

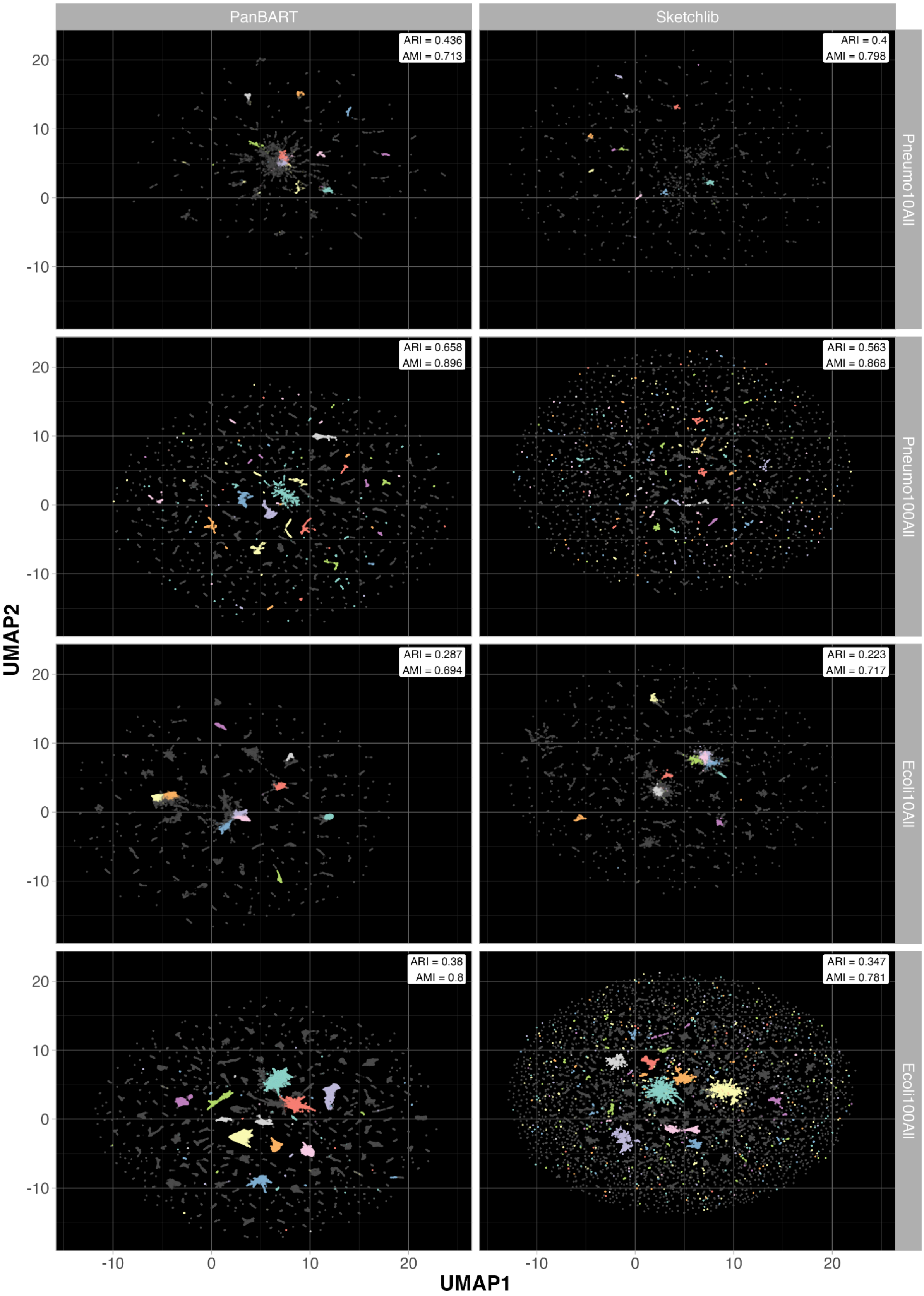

Supplementary Figure 12: Leiden cluster assignment using embedding space of PanBART and Sketchlib for training data for models trained on all lineages. Adjusted Rand Index (ARI) and Adjusted Mutual Information (AMI) between assigned clusters and PopPUNK clusters are displayed in the top right of each plot. Each point represents a single genome, the top 10 largest clusters are coloured.

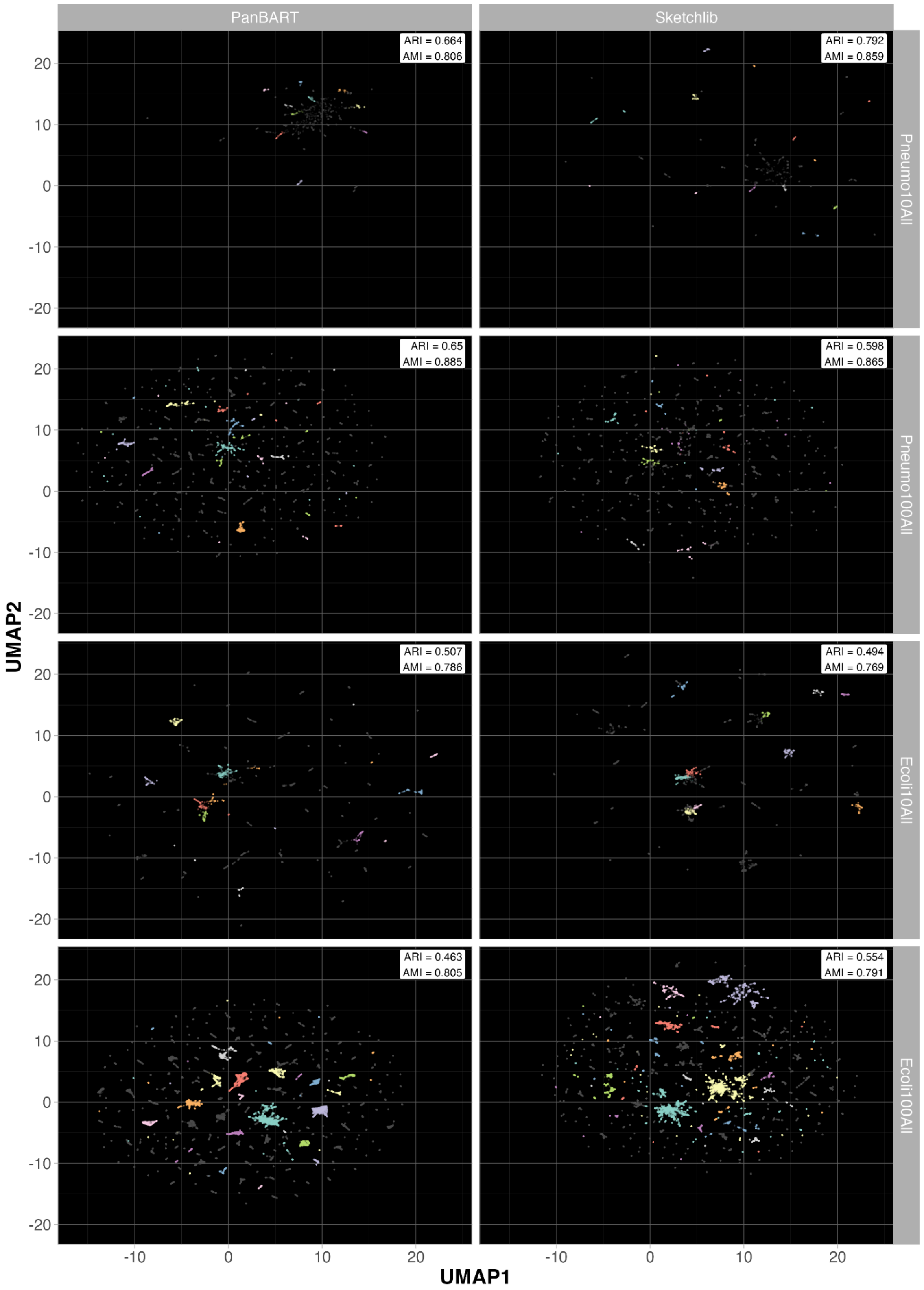

Supplementary Figure 13: Leiden cluster assignment using embedding space of PanBART and Sketchlib for testing data including all lineages. Adjusted Rand Index (ARI) and Adjusted Mutual Information (AMI) between assigned clusters and PopPUNK clusters are displayed in the top right of each plot. Each point represents a single genome, the top 10 largest clusters are coloured.

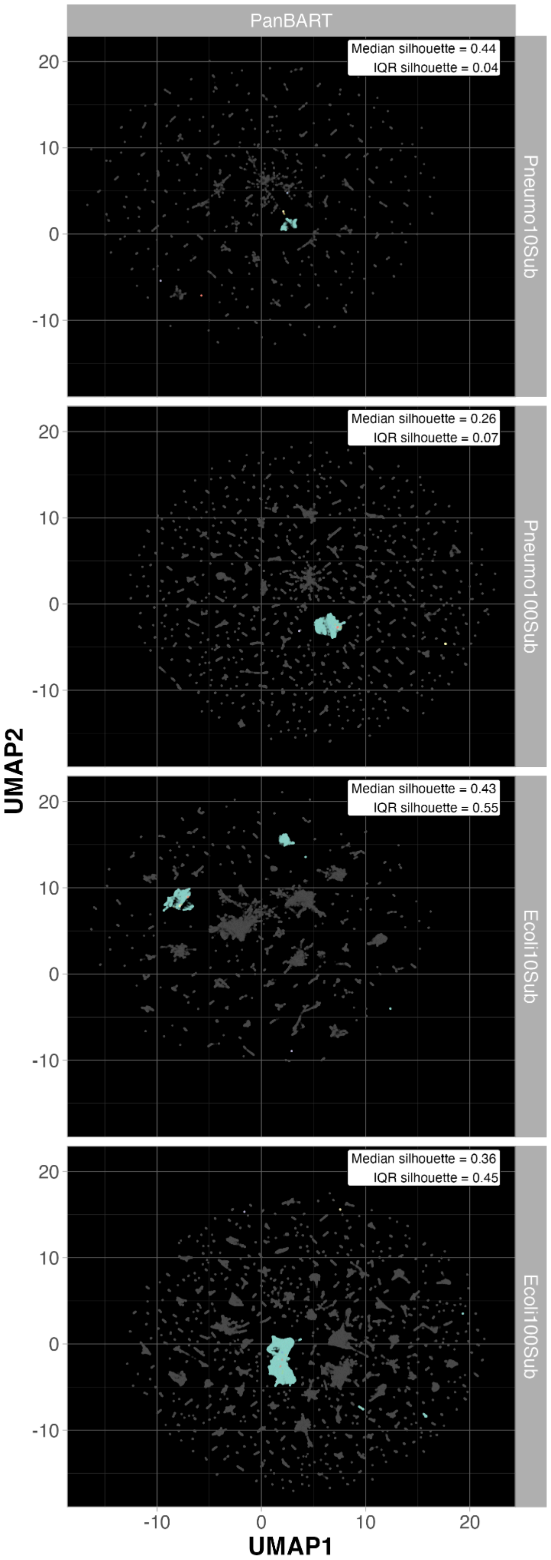

Supplementary Figure 14: PanBART genome embeddings for novel held-out lineages data (coloured) overlaid onto training data (grey), where the model was training on a subset of lineages. Each point represents the embedding of a single genome. Colours show the 10 largest PopPUNK held-out lineage clusters for each species. The median and Inter-Quartile Range (IQR) of silhouette scores for the 10 largest PopPUNK lineage clusters is shown in the top right corner of each plot.

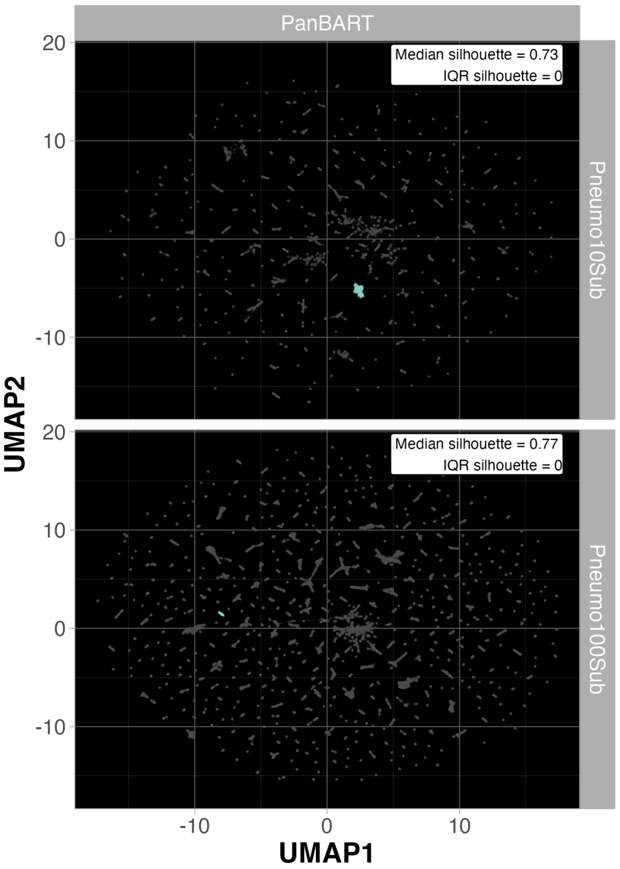

Supplementary Figure 15: PanBART genome embeddings for *S. mitis* data (coloured) overlaid onto training data (grey), where the model was training on a subset of lineages of *S. pneumoniae*. Each point represents the embedding of a single genome. Colours show only *S. mitis* genomes. The median and Inter-Quartile Range (IQR) of silhouette scores for the 10 largest PopPUNK lineage clusters is shown in the top right corner of each plot.

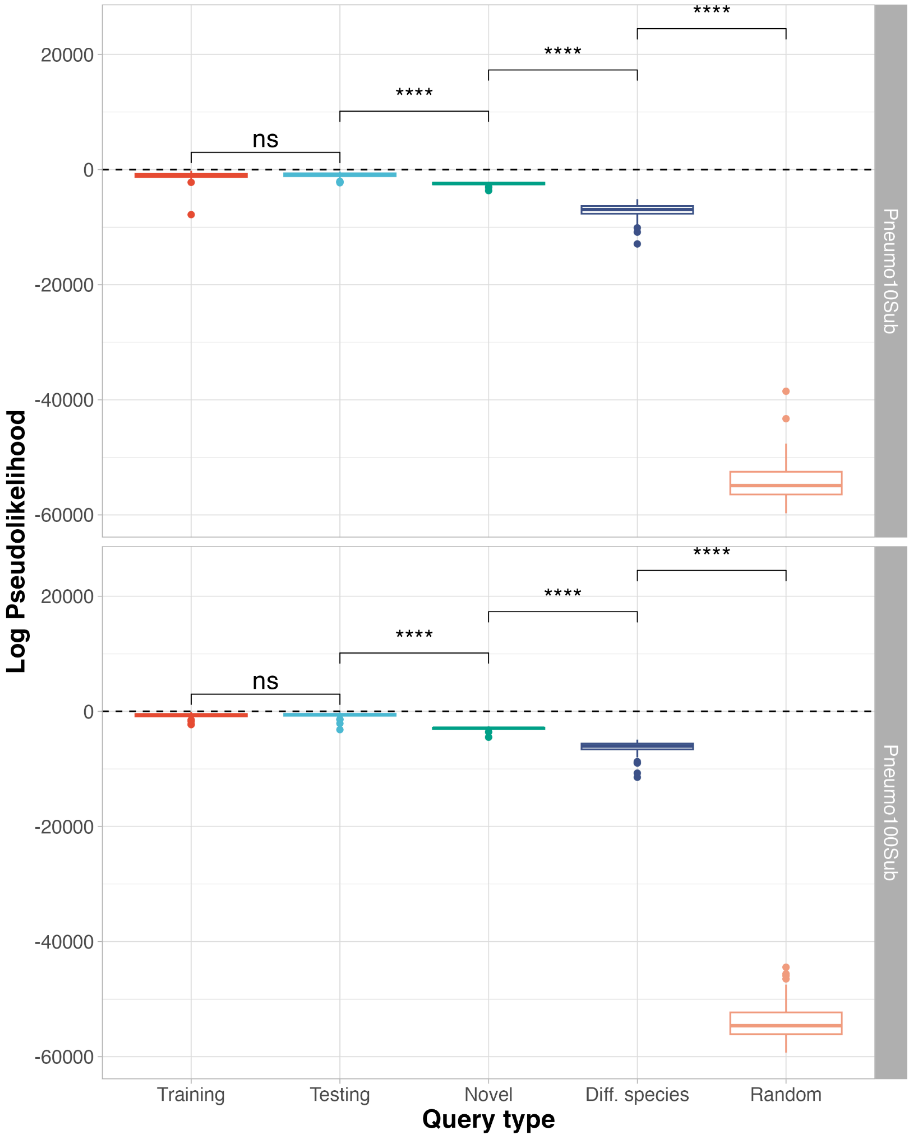

Supplementary Figure 16: Differentiation of observed and novel genomes by PanBART. Query types: “Training”, genomes seen by the model in training; “Testing”, genomes belonging to PopPUNK lineages used in training, but the genomes themselves were not seen in training; “Novel”, genomes belonging to PopPUNK lineages not used in training; “Diff. species”, genomes belonging to a closely related Streptococcal species, *Streptococcus mitis*, which was not used in training; “Random”, training genomes with gene order randomised before passing to PanBART. Each query type contains 50 genomes. Statistical comparisons conducted with Wilcoxon test: ns, $p > 0.05$; ****, $p \leq 0.0001$.

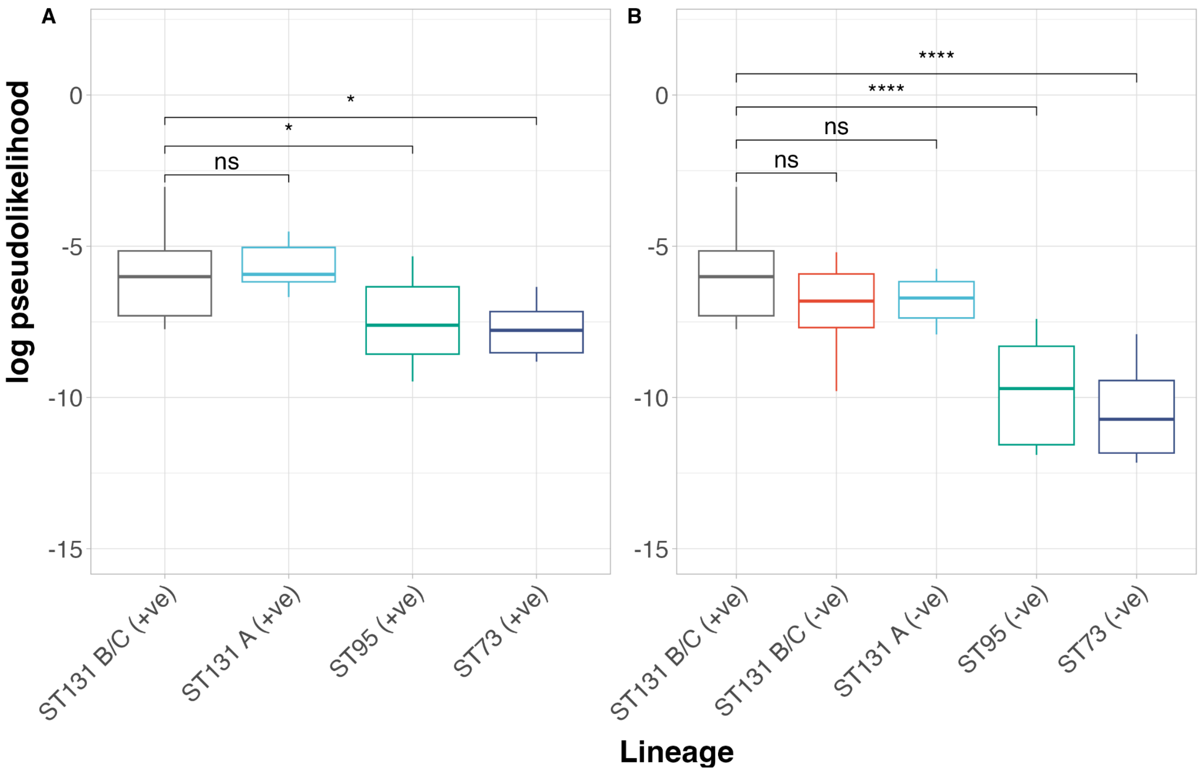

Supplementary Figure 17: Log pseudolikelihood comparison of *blaCTX-M* uptake potential across *E. coli* lineages. Analysis was split between genomes which contained *blaCTX-M* but had it bioinformatically removed prior to analysis (labelled "+ve") (**A**) and genomes which did not contain *blaCTX-M* (labelled "-ve") (**B**). Comparisons conducted using genomes used in PanBART training. Each boxplot represents 10 genomes. ST131 B/C (+ve) was shared between both analyses as a positive control for statistical comparisons. Statistical comparisons conducted with Wilcoxon test: ns, p >0.05; *, p ≤ 0.05; ***, p ≤ 0.001; ****, p ≤ 0.0001.

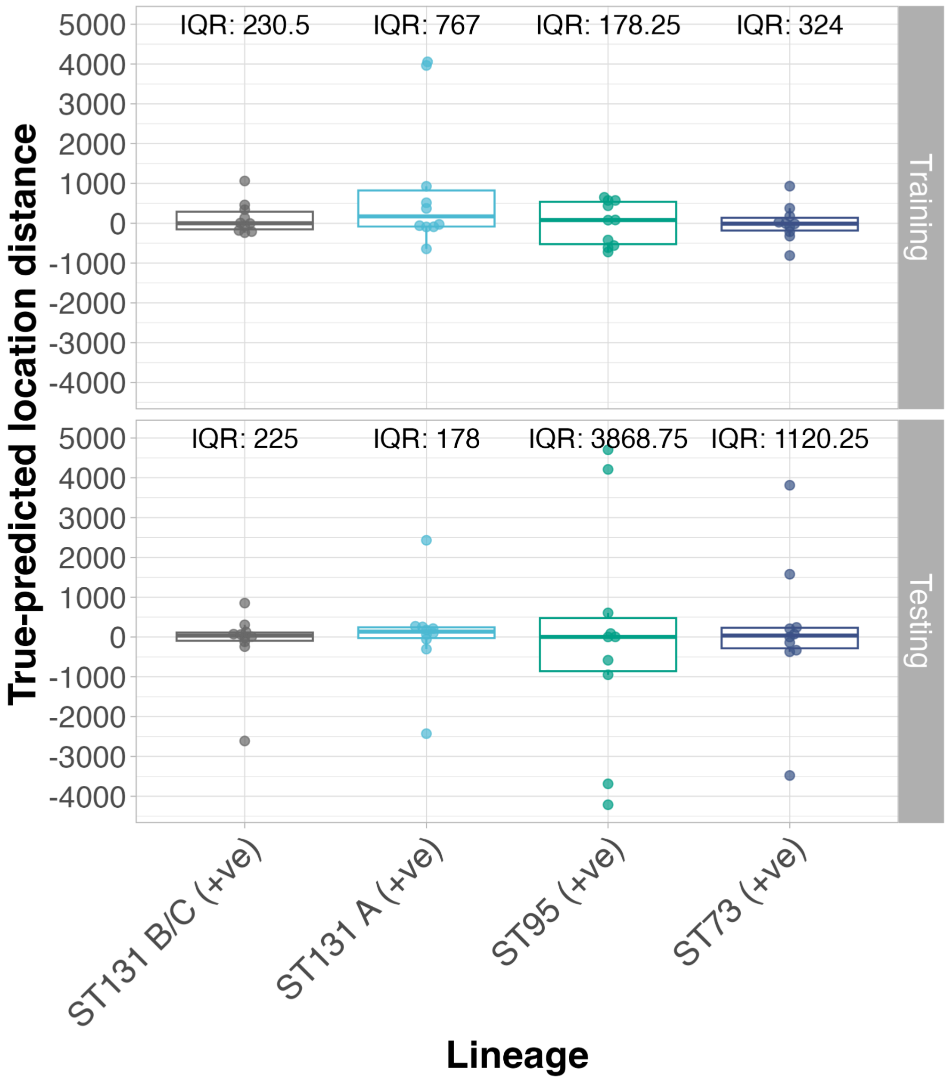

Supplementary Figure 18: Difference between true and predicted position of *blaCTX-M* in *E.coli* genomes naturally containing *blaCTX-M*. Inter-quartile range (IQR) shown for each dataset and lineage analysed.

### Supplementary Tables

Supplementary Table 1: Summary of genome counts used in model training and analysis.

| Species model | Type | Data | Lineages | Count |
|---|---|---|---|---|
| *E. coli* | Novel | 10% Data | Subset | 3609 |
| *E. coli* | Novel | 100% Data | Subset | 35791 |
| *E. coli* | Testing | 10% Data | All | 3568 |
| *E. coli* | Testing | 10% Data | Subset | 3212 |
| *E. coli* | Testing | 100% Data | All | 38168 |
| *E. coli* | Testing | 100% Data | Subset | 34593 |
| *E. coli* | Training | 10% Data | All | 40063 |
| *E. coli* | Training | 10% Data | Subset | 37173 |
| *E. coli* | Training | 100% Data | All | 316024 |
| *E. coli* | Training | 100% Data | Subset | 287388 |
| *E. coli* | Validation | 10% Data | All | 5278 |
| *E. coli* | Validation | 10% Data | Subset | 4915 |
| *E. coli* | Validation | 100% Data | All | 39546 |
| *E. coli* | Validation | 100% Data | Subset | 35966 |
| *S. pneumoniae* | Novel | 10% Data | Subset | 635 |
| *S. pneumoniae* | Novel | 100% Data | Subset | 6034 |
| *S. pneumoniae* | S. mitis | 10% Data | All | 399 |
| *S. pneumoniae* | S. mitis | 100% Data | All | 399 |
| *S. pneumoniae* | Testing | 10% Data | All | 998 |
| *S. pneumoniae* | Testing | 10% Data | Subset | 939 |
| *S. pneumoniae* | Testing | 100% Data | All | 11358 |
| *S. pneumoniae* | Testing | 100% Data | Subset | 10758 |
| *S. pneumoniae* | Training | 10% Data | All | 13419 |
| *S. pneumoniae* | Training | 10% Data | Subset | 12908 |
| *S. pneumoniae* | Training | 100% Data | All | 96020 |
| *S. pneumoniae* | Training | 100% Data | Subset | 91190 |
| *S. pneumoniae* | Validation | 10% Data | All | 1834 |
| *S. pneumoniae* | Validation | 10% Data | Subset | 1769 |
| *S. pneumoniae* | Validation | 100% Data | All | 12054 |
| *S. pneumoniae* | Validation | 100% Data | Subset | 11450 |

Supplementary Table 2: Dataset nomenclature and model sizes.

| Species model | Data | Lineages | Training genome count | Name | No. model parameters | No. GPUs for training | Training time (hours) |
|---|---|---|---|---|---|---|---|
| *S. pneumoniae* | 10% Data | Subset | 12908 | Pneumo10Sub | 28,084,480 | 4 | 190 |
| *S. pneumoniae* | 100% Data | Subset | 91190 | Pneumo100Sub | 28,563,200 | 4 | 574 |
| *S. pneumoniae* | 10% Data | All | 13419 | Pneumo10All | 28,040,960 | 4 | 262 |
| *S. pneumoniae* | 100% Data | All | 96020 | Pneumo100All | 28,563,200 | 4 | 585 |
| *E. coli* | 10% Data | Subset | 37173 | Ecoli10Sub | 51,562,240 | 4 | 564 |
| *E. coli* | 100% Data | Subset | 287388 | Ecoli100Sub | 51,688,192 | 4 | 599 |
| *E. coli* | 10% Data | All | 40063 | Ecoli10All | 51,715,840 | 4 | 547 |
| *E. coli* | 100% Data | All | 316024 | Ecoli100All | 51,743,488 | 4 | 632 |

Supplementary Table 3: Speed of testing data lineage assignment. Sketchlib runtime calculated as sum of sketching and distance calculations.

| Dataset Name | No. Genomes | PanBART runtime (mins) | PanBART genomes/min | Sketchlib runtime (mins) | Sketchlib genomes/min |
|---|---|---|---|---|---|
| Pneumo10Sub | 939 | 5.43 | 173 | 0.43 | 2209 |
| Pneumo100Sub | 10758 | 63.05 | 171 | 4.87 | 2208 |
| Pneumo10All | 998 | 5.80 | 172 | 0.45 | 2211 |
| Pneumo100All | 11358 | 66.92 | 170 | 5.13 | 2213 |
| Ecoli10Sub | 3212 | 28.27 | 114 | 2.67 | 1201 |
| Ecoli100Sub | 34593 | 309.93 | 112 | 28.66 | 1207 |
| Ecoli10All | 3568 | 31.73 | 112 | 2.96 | 1204 |
| Ecoli100All | 38168 | 339.09 | 113 | 31.75 | 1202 |

Supplementary Table 4: Frequency of *blaCTX-M* in *E. coli* genomes in AllTheBacteria dataset.

| Lineage | ***blaCTX-M*** **positive** | ***blaCTX-M*** **negative** | Total count | % positive |
|---|---|---|---|---|
| ST73 | 281 | 6723 | 7004 | 4.01 |
| ST95 | 321 | 4928 | 5249 | 6.12 |
| ST131 A | 2114 | 1015 | 3129 | 67.56 |
| ST131 B/C | 11210 | 4451 | 15661 | 71.57 |

Supplementary Table 5: Functional Annotation of extreme SHAP outliers associated with *clbB* gene.

| **SHAP** | **Locus** | **Gene** | **Product** | **UniRef** | **Peak** |
|---|---|---|---|---|---|
| -0.011412503 | 2144 | tsaB | tRNA threonylcarbamoyladenosine biosynthesis protein TsaB | UniRef50_A0A7H4MWN0 | 1 |
| | 2145 | yoaB | RutC family protein YoaB | UniRef50_P0AEB8 | |
| | 2146 | NA | hypothetical protein | NA | |
| | 2147 | NA | hypothetical protein | NA | |
| | 2148 | NA | hypothetical protein | NA | |
| | 2149 | NA | hypothetical protein | NA | |
| | 2150 | NA | hypothetical protein | NA | |
| | 2151 | NA | hypothetical protein | NA | |
| | 2152 | yoaE | CNNM family cation transport protein YoaE | UniRef50_P0AEC2, UniRef90_UPI000541D78E | |
| | 2153 | yoaL | Protein YoaL | UniRef50_P0DPP2, UniRef90_P0DPP2 | |
| | 2154 | manX | PTS mannose transporter subunit IIAB | UniRef50_P69799, UniRef90_A0A0F1R454 | |
| | 2155 | NA | hypothetical protein | NA | |
| | 2156 | NA | hypothetical protein | NA | |
| | 2157 | NA | hypothetical protein | NA | |
| | 2158 | mntP | manganese efflux pump MntP | UniRef50_Q3Z2G2, UniRef90_Q0T515 | |
| | 2159 | rlmA | 23S rRNA (guanine(745)-N(1))-methyltransferase | UniRef50_A0A0D7LXL9, UniRef90_A0A0K4LGX5 | |
| | 2160 | NA | hypothetical protein | NA | |
| | 2161 | NA | hypothetical protein | NA | |
| | 2162 | NA | hypothetical protein | NA | |
| | 2163 | kdgR | DNA-binding transcriptional regulator KdgR | UniRef50_P37728, UniRef90_A0A0A5P595 | |
| | 2164 | NA | hypothetical protein | NA | |
| | 2165 | htpX | protease HtpX | UniRef50_Q6D4G3, UniRef90_A8AFM2 | |
| | 2166 | prc | carboxy terminal-processing peptidase | UniRef50_P23865, UniRef90_A0A090V0F9 | |
| | 2167 | proQ | RNA chaperone ProQ | UniRef50_Q5E5C2, UniRef90_A0A2A6QDW9 | |
| | 2168 | msrC | Free methionine-R-sulfoxide reductase | UniRef50_P76270, UniRef90_P76270 | |
| | 2169 | yebS | membrane integrity lipid transport subunit YebS | UniRef50_P0AD04 | |

| | | | | | |
|---|---|---|---|---|---|
| | 2170 | NA | hypothetical protein | NA | |
| | 2171 | NA | Ribosomal RNA small subunit methyltransferase F | UniRef50_Q8XCL9 | |
| | 2172 | NA | hypothetical protein | NA | |
| | 2173 | NA | hypothetical protein | NA | |
| | 2174 | NA | Serine/threonine-protein phosphatase 1 | UniRef50_Q8ZNY9 | |
| | 2175 | NA | DUF2511 domain-containing protein | UniRef50_A0A4U9HY52 | |
| -0.008460756 | 4144 | NA | hypothetical protein | NA | **2** |
| | 4145 | NA | hypothetical protein | NA | |
| | 4146 | NA | hypothetical protein | NA | |
| | 4147 | NA | transposase | UniRef50_A0A8T5Z4I5, UniRef90_UPI0010B78271 | |
| | 4148 | NA | hypothetical protein | UniRef50_A0A3P5DKH8, UniRef90_A0A3P5DKH8 | |
| | 4149 | NA | RanBP2-type domain-containing protein | UniRef50_A0A7Z1QA92 | |
| | 4150 | NA | hypothetical protein | NA | |
| | 4151 | NA | hypothetical protein | NA | |
| | 4152 | NA | hypothetical protein | NA | |
| | 4153 | NA | hypothetical protein | NA | |
| | 4154 | cbtA | Toxin CbtA | UniRef50_P64525 | |
| | 4155 | NA | hypothetical protein | NA | |
| | 4156 | NA | DUF957 domain-containing protein | UniRef50_Q5K5L1, UniRef90_Q707E9 | |
| | 4157 | NA | Restriction methylase | UniRef50_B7LDP3 | |
| | 4158 | NA | Nitrite extrusion protein 2 | UniRef50_I6CVP5 | |
| | 4159 | NA | HNH endonuclease domain protein | UniRef50_A0A090VNZ7, UniRef90_V0ZCE9 | |
| 0.007657335 | 4352 | gloB | hydroxyacylglutathione hydrolase | UniRef50_P0AC85, UniRef90_P0AC85 | **3** |
| | 4353 | yafS | Uncharacterized protein YafS | UniRef50_P75672, UniRef90_P75672 | |
| | 4354 | NA | Ribonuclease HI | UniRef50_P43807 | |
| | 4355 | dnaQ | DNA polymerase III subunit epsilon | UniRef50_P03007, UniRef90_A0A2X3M5V0 | |
| | 4356 | NA | hypothetical protein | NA | |
| | 4357 | NA | hypothetical protein | NA | |
| | 4358 | NA | hypothetical protein | NA | |

| | | | | | |
|---|---|---|---|---|---|
| | 4359 | NA | hypothetical protein | NA | |
| | 4360 | impA | ImpA domain protein | UniRef50_W8STY3 | |
| | 4361 | tssM | type VI secretion system membrane subunit TssM | UniRef50_X5FDK4, UniRef90_UPI001F3F3BDB | |
| | 4362 | NA | Protein rhsD, truncation | UniRef50_A0A1L4IZ17 | |
| | 4363 | NA | Lipoprotein | UniRef50_A0A0B1JN69 | |
| | 4365 | NA | Hydrolase | UniRef50_A0A0H2VAN3, UniRef90_A0A8E0IVD5 | |
| | 4366 | NA | hypothetical protein | NA | |
| | 4367 | NA | Sialate O-acetylesterase domain-containing protein | UniRef50_A0A2Z2J3G7, UniRef90_A0A2K0PKG1 | |
| 0.008413226 | 4368 | NA | hypothetical protein | NA | 4 |
| | 4369 | iucA | aerobactin synthase IucA | NA | |
| | 4370 | iucB | N(6)-hydroxylysine O-acetyltransferase IucB | UniRef50_Q47317, UniRef90_Q47317 | |
| | 4371 | iucA | aerobactin synthase IucA | NA | |
| | 4372 | iucC | NIS family aerobactin synthetase IucC | NA | |
| | 4373 | iutA | ferric aerobactin receptor IutA | UniRef50_P14542, UniRef90_A0A379ZE27 | |
| | 4374 | sat | serine protease autotransporter toxin Sat | UniRef50_Q8FDW4, UniRef90_Q8FDW4 | |
| | 4375 | NA | hypothetical protein | UniRef50_A0A0H2VC54, UniRef90_A0A0H2VC54 | |
| | 4377 | yhiS | YhiS | UniRef50_Q8X5P7, UniRef90_F5P2P9 | |
| | 4378 | arsC | glutaredoxin-dependent arsenate reductase | UniRef50_P08692, UniRef90_A0A5Z7L310 | |
| | 4379 | arsR | As(III)-sensing metalloregulatory transcriptional repressor ArsR | UniRef50_P30329, UniRef90_P0AB94 | |
| | 4380 | arsR | As(III)-sensing metalloregulatory transcriptional repressor ArsR | UniRef50_P15905 | |
| | 4381 | NA | hypothetical protein | NA | |
| | 4382 | gorA | glutathione-disulfide reductase | UniRef50_A0A2K5XF66, UniRef90_UPI0005A68B79 | |
| | 4383 | rlmJ | Ribosomal RNA large subunit methyltransferase J | UniRef50_P37634, UniRef90_P37634 | |
| 0.012660529 | 4384 | NA | hypothetical protein | NA | 5 |
| | 4385 | NA | Ribosomal RNA small subunit methyltransferase J | UniRef50_Q1R5B9 | |
| | 4386 | dtpB | dipeptide/tripeptide permease DtpB | UniRef50_P36837, UniRef90_P36837 | |
| | 4387 | NA | hypothetical protein | NA | |

| | | | | |
|---|---|---|---|---|
| | 4388 | NA | hypothetical protein | NA |
| | 4389 | NA | hypothetical protein | NA |
| | 4390 | NA | hypothetical protein | NA |
| | 4391 | NA | hypothetical protein | NA |
| | 4394 | NA | Zinc ribbon domain-containing protein | UniRef50_H5V7N2, UniRef90_H5V7N2 |
| | 4395 | NA | Phage head-tail connector protein | UniRef50_H5V7N1, UniRef90_H5V7N1 |
| | 4396 | tnp | IS1 family transposase | UniRef100_UPI000450A93A, UniRef50_A0A658YYP2, UniRef90_A0A827ZTK5 |
| | 4397 | NA | IS1 transposase | UniRef50_A0A0A6ZQ01, UniRef90_A0A2S1JA09 |
| | 4399 | lutC | L-lactate utilization protein LutC, contains LUD domain | UniRef50_P77433, UniRef90_D4BEN2 |

Supplementary Table 6: Species with >5000 genomes in AllTheBacteria dataset (release 0.2, increment 2024.08). Species assignments from GTDB (Chaumeil et al. 2022).

| **Species** | **Genome count** |
|---|---|
| *Salmonella enterica* | 645471 |
| *Escherichia coli* | 398298 |
| *Mycobacterium tuberculosis* | 183795 |
| *Streptococcus pneumoniae* | 122512 |
| *Staphylococcus aureus* | 122452 |
| *Campylobacter_D jejuni* | 95357 |
| *Klebsiella pneumoniae* | 74096 |
| *Neisseria gonorrhoeae* | 57716 |
| *Streptococcus pyogenes* | 46669 |
| *Pseudomonas aeruginosa* | 43732 |
| *Streptococcus agalactiae* | 37669 |
| *Enterococcus_B faecium* | 36841 |
| *Neisseria meningitidis* | 36101 |
| *Campylobacter_D coli* | 34890 |
| *Listeria monocytogenes* | 32676 |
| *Listeria monocytogenes_B* | 32437 |
| *Clostridioides difficile* | 29433 |
| *Acinetobacter baumannii* | 28645 |
| *Vibrio cholerae* | 13240 |
| *Enterococcus faecalis* | 9885 |
| *Haemophilus influenzae_E* | 9429 |
| *Vibrio parahaemolyticus* | 8342 |
| *Bordetella pertussis* | 7522 |
| *Enterobacter hormaechei_C* | 7486 |
| *Legionella pneumophila* | 7055 |
| *Staphylococcus epidermidis* | 6591 |
| *Burkholderia mallei* | 6461 |
| *Mycobacterium abscessus* | 6140 |
| *Staphylococcus pseudintermedius* | 5534 |
